# From Fabrication to Flow: Impact of Print Orientation on Surface Qualities and Capillary-Driven Flow in Laser SLA-based Open Microchannels

**DOI:** 10.64898/2026.04.10.717746

**Authors:** Ariel Lin, Laura A. Milton, Damon Wing Hey Chan, Nidhi Ghadge, Jodie C. Tokihiro, Lauren G. Brown, Albert Shin, Yi-Chin Toh, Ayokunle O. Olanrewaju, Erwin Berthier, Jean Berthier, Ashleigh B. Theberge

## Abstract

Stereolithography (SLA) 3D printing has become increasingly popular for fabricating microfluidic devices, with applications including hydrogel patterning and tissue modeling. In open-channel systems with surface tension-driven flow, 3D-printer-induced discrepancies in channel surface texture can significantly impact fluid flow and device performance. While previous work has focused on comparing different 3D printing methods for microchannel fabrication, the effect of device orientation during SLA printing on microchannel morphology and capillary-driven flow has not been systematically evaluated. Furthermore, there is minimal research elucidating the influence of channel surface texture on the flow of biologically relevant hydrogel precursors commonly used in organ-on-a-chip applications. Herein, we investigated the impact of print orientation on channel morphology, fluid wetting behavior, and fluid flow by comparing laser SLA-based parts where the length of the channel was tilted at 0°, 15°, 45°, or 90° during printing. We demonstrated that channel floor surface texture is greatly affected by print orientation: the highest axial surface roughness was measured in 15° printed channels, and the highest axial surface tortuosity–which describes the real length along the surface–was measured in 45° printed channels. Print angles of 15° and 45° also resulted in asymmetric roughness of the channel floor, which caused asymmetric wetting of glycerol solution. Surface tension-driven flow of glycerol solution, agarose precursor solution, and collagen precursor solution was affected by print orientation, in which the 45° printed flow devices had slowest flow for all test fluids. Root mean square roughness was not a reliable predictor of slower flow; instead, surface tortuosity should be considered. Potential alternatives to better theoretically model how print angle-induced surface texture affects open-channel flow are discussed as well. These findings provide a framework of fabrication considerations for laser SLA printing of open microchannels that can also be applied to other layer-by-layer, vat photopolymerization-based 3D printing technologies.

## INTRODUCTION

Microfluidic technologies, which enable experiments on the nanometer to micrometer scale, have been adopted across many scientific disciplines with advantages such as decreased sample consumption, increased throughput, and lower cost.^1^ Historically, microfluidic devices were mainly fabricated through soft lithography and replica molding with polydimethylsiloxane (PDMS). PDMS has certain advantages as a transparent and largely biocompatible material;^2,3^ however, its associated fabrication technique, soft lithography, is time and labor intensive.^4^ In addition, more complex microfluidic devices often require multiple layers of PDMS to be molded, aligned, and bonded together through a manual process that is heavily dependent on user skill and prone to poor reproducibility.^5^ To overcome the limitations of soft lithography, scientists have pivoted towards other methods including computer numerical control milling^6^ and 3D printing,^7–12^ which allow for direct device fabrication without the need for master templates.

3D printing is an additive manufacturing process that creates structures layer by layer, usually from polymeric materials.^13^ For microfluidic device fabrication, the most commonly used techniques are fused filament fabrication (FFF) and vat photopolymerization.^7–9^ FFF employs a nozzle extruding melted polymer filaments to build each layer. This printing method is low-cost but has less precision than other methods, and greater surface texture is generated on these devices.^14^ Vat photopolymerization, which includes laser stereolithography (SLA), digital light processing SLA (DLP-SLA), and liquid crystal display SLA (LCD-SLA), involves selective curing of each layer in a photopolymerizable resin bath. Vat photopolymerization enables higher print resolution and better surface finish but requires additional post-processing steps to ensure the removal of unreacted material.^10,12,15–18^ SLA (laser, DLP, and LCD) printing has become one of the most commonly used methods for microfluidic fabrication. Gonzalez et al.^9^ showed that between 2008 and 2022, more studies with the keyword ‘SLA’ were published compared to FFF and PDMS-based methods for the fabrication of microfluidic devices.

Besides the transition to direct fabrication methods over the past decade, many microfluidicists have also adopted open microfluidic designs where at least one of the microfluidic channel walls is removed to introduce an air-liquid interface. Open microfluidics provides several advantages over traditional closed-channel (i.e., a channel completely enclosed by four walls) counterparts, including affordability, convenience of fabrication, and ease of channel surface modification.^19^ Moreover, open microchannels provide the ability to easily add, remove, and interact with the sample at any point in the channel. Open-channel systems have been extensively used to create 3D cell culture models that mimic tissues and organs, where the channels are leveraged to pattern, or shape, cell-laden hydrogel precursor solutions such as collagen or fibrin.^20–26^ Hydrogel precursors are considered to be complex fluids and have physical properties that differ from fluids conventionally used in studies of microfluidic flow dynamics (e.g., water, nonanol). Namely, these hydrogel precursor fluids are mildly shear thinning–the viscosity decreases as shear rate increases. Thus, hydrogel precursors are expected to have different flow profiles from traditionally used test fluids. Understanding how the flow of hydrogel precursors is influenced by the microfluidic channel surface is critical to developing effective hydrogel patterning platforms.

In many open microchannel designs, flow is driven by spontaneous capillary flow (i.e., surface tension forces) and does not require external actuation or pumping.^19,27–30^ Hence, the interaction between the fluid and the surface of the channel significantly affects the fluid flow and performance of the microfluidic device. Previous work has shown that FFF-based microchannels possess significant surface roughness that affects flow behavior, and the induced surface texture is dependent on part orientation during the printing process.^31,32^ Li et al.^31^ showed that FFF printing at different *xy*-orientations imparts different surface texture patterns that could be leveraged to tune microfluidic mixing in a closed channel system. Lade et al.^32^ found that the *xy*-orientation during FFF printing induces varying surface roughness, and that roughness causes the fluid front to move in a pulsing motion down the channel due to contact line pinning; surface roughness and fluid flow in laser SLA-based microchannels was also explored, but only one print orientation was tested. Due to the popularity of SLA printing amongst microfluidic users, further research on the effects of SLA print orientation on microchannel fabrication and fluid flow is warranted.

In this work, we assessed the effect of *z*-orientation on laser SLA-based microchannels by printing shallow rectangular open microchannels with the length of the channel angled at 0°, 15°, 45°, or 90°. The impact of print orientation on channel morphology and fluid spreading was evaluated with scanning electron microscopy (SEM), profilometry, and goniometry. Capillary driven flow with the laser SLA-based microchannels printed at different angles was also tested with glycerol solution (an aqueous Newtonian fluid) as well as agarose and collagen precursor solutions (biologically relevant, complex fluids). We found that channel orientation during printing greatly impacts the surface texture of the microchannel floors and moderately impacts the channel walls. The resulting texture affects the spreading behavior of test fluids on the surface and therefore also affects capillary-driven flow. Channel filling was slowest in devices printed at 45°, but root mean square roughness (R_q_)–a commonly used parameter to report surface roughness– did not account for the delayed velocity. Instead, we propose the introduction of two parameters to better approximate the effect of surface roughness on capillary-driven flow in open channels: (1) apparent average friction length inclusive of axial surface tortuosity and (2) average apparent contact angle. The first parameter pertains to the rough wall friction along the length of the channel. The second parameter concerns the capillary pressure of the advancing fluid front, which fluctuates when the three-phase contact line passes over roughness reliefs.

## EXPERIMENTAL SECTION

### Microchannel Design and Fabrication

Flow devices were designed to have two circular reservoirs connected by a rectangular open channel with rounded bottom edges. The designed channel dimensions varied to achieve within 5% error of width=2 mm width height=0.8 mm (Table SI.3.1). The channel dimensions were verified using a Keyence wide area 3D measurement VR-5000 profilometer (Keyence Corporation of America, Itasca, Illinois) after applying a 3D laser scanning spray (Helling, Heidgraben, Germany) to reduce the opacity of the devices. Profilometry blocks, used for SEM and stylus profilometry, were designed with two full channels flanked by two half channels. The full channels had cross section dimensions of width=1mm and height=2 mm. Contact angle blocks, used for goniometry, were designed to mimic the surface of the channel floor and wall. Images and schematics of the device and block designs can be found in Figure 1A-C. Engineering drawings can be found in (Fig. SI.3.1).

**Figure 1.**
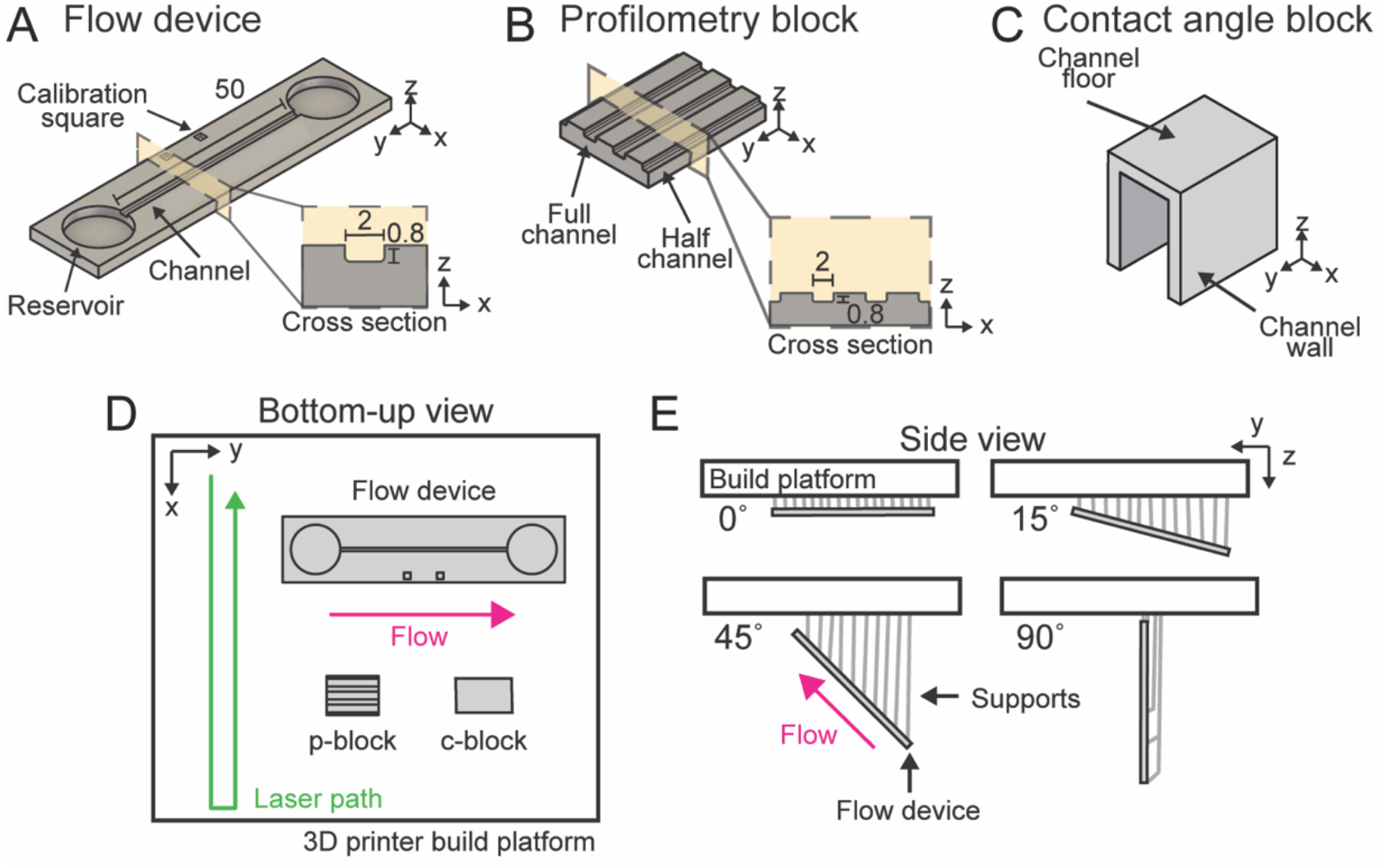
Schematic of microchannels and part orientation during printing. (A) The flow device has a shallow rectangular channel with rounded corners. Cross section shows slice in the *z-x* plane. (B) The profilometry block has two full channels with the same height/width dimensions as the flow device and two flanking half channels. Cross section shows slice in the *z-x* plane. Units for (A) and (B) are in mm. (C) The contact angle block has multiple sides to mimic the channel floor and wall with respect to print orientation. (D) Bottom-up view of microchannels during printing. Parts are oriented on the build platform with the channel(s) perpendicular to the laser path. (E) Side-view of microchannels during printing. Parts are printed upside down with the channel floor facing away from the supports.

All designs were made using Autodesk Fusion 360 (Autodesk, San Francisco, California) or Solidworks (SolidWorks, Waltham, Massachusetts) and exported as Standard Tessellation Language (STL) files. STL files were imported into PreForm (Formlabs, Somerville, Massachusetts) to prepare models with 0.1 mm *z*-slices. Parts were orientated with the channels perpendicular to the laser path and tilted at 0°, 15°, 45°, and 90° along the length of the channel (Fig. 1D and E). Parts were printed with Formlabs clear resin V4 on a FormLabs 3B stereolithography printer. After the printing, the devices were washed with three consecutive isopropanol alcohol washes for a total of one hour, dried with compressed air, and placed in a Formcure to postcure for 30 min at 25°C.

### Microchannel Surface Texture and Wetting Characterization

Image capture of the profilometry block microchannels was performed using the Apreo-2S SEM (ThermoFisher Scientific, Waltham, Massachusetts) after coating the blocks with 4.5 to 5 nm of platinum using a Leica EM ACE600 sputter coater (Leica Microsystems, Wetzlar, Germany). Surface topography measurements of profilometry blocks and contact angle blocks were conducted using a Bruker OM-DektakXT stylus profilometer (Bruker, Billerica, MA).

Measurements were taken over a scan length of 3 mm and 2-3 mm for the profilometry and contact angle blocks, respectively. A Gaussian regression filter was applied with a short cutoff of 2.5 µm and a long cutoff of 0.8 mm to obtain the roughness and waviness profiles. Each of the plotted R_q_ (root mean square roughness) and RSm (mean spacing between roughness profile irregularities) data points represents an average of at least three measurements on an independently printed sample. Reported tortuosity values represent an average of four measurements performed on three independently printed samples.

Contact angle measurements on contact angle blocks were conducted with a Kruss DSA-25E drop shape analyzer (Kruss GmbH, Hamburg, Germany). Measurements were taken with a window of 5 to15 minutes after oxygen-plasma treatment with a Diener Zepto PC EX Type PB plasma treater (Diener electronic GmbH & Co. KG, Ebhausen, Germany). Each of the plotted contact angle data points represent the average of three sessile drop measurements on a single 2 µL glycerol solution droplet collected with the Kruss ADVANCE software.

### Solution Preparation and Physical Properties

0.644 mass fraction glycerol (Fisher Scientific, Grand Island, New York) was prepared by diluting in ASTM H2O (MilliporeSigma, Darmstadt, Delaware). The glycerol solution was kept at room temperature (16-21° C) for all experiments. 1.5% (w/v) agarose precursor solution was prepared by dissolving low-temperature gelling agarose (Sigma Aldrich, St. Louis, Missouri), in ASTM H2O in a microwave. Agarose precursor solution was kept at room temperature when not in use, and heated to 60°C until use for all experiments. A 3 mg/mL collagen precursor solution was prepared by neutralizing concentrated rat tail collagen (Corning, Harrodsburg, Kentucky) with phosphate buffer saline (Gibco, Grand Island, New York), HEPES (Corning), sodium bicarbonate (Corning), sodium hydroxide (Fisher Scientific), and ASTM H2O. Collagen precursor solution was prepared within 10 minutes of the experiment and kept on ice until use. All solutions were colored with 1.2% (v/v) blue food coloring (McCormick, Hunt Valley, Maryland).

The surface tension of the test fluids was measured with a Kruss DSA-25E drop shape analyzer using the pendant drop method on the Kruss ADVANCE software. Each value represents the average of three independently performed drops. For contact angle measurements, glass slides coated with approximately 1 mL of clear resin were fabricated to obtain smooth test surfaces. The contact angle of the test fluids was measured as described above with 8 µL droplets. The droplet volume was increased to prevent quick gelling of the agarose precursor solution. Measurements were taken within a window of 5-10 minutes after plasma treatment. The viscosity of the solutions was measured using a Discovery HR-2 rheometer (TA instruments, New Castle, Delaware) with a 40 mm one degree cone geometry. Depending on the test fluid, 308-350 µL of samples were loaded. Initial experiments were performed to determine the effective pre-shear and equilibration conditions for each test fluid. Shear sweep experiments from 0.5 to 500 s-1 were performed at 19°C, 60°C, and 8°C for glycerol solution, agarose precursor, and collagen precursor, respectively.

### Open-channel Flow Experiments

Flow devices were plasma treated 5-15 minutes prior to use and positioned on a leveled white stage with an adjustable lab jack. Tabletop studio lights were adjusted to minimize shadows and maximize visibility of the fluid front. The fill volume of the fluid reservoir ranged from 455-490 µL depending on the print orientation of the device and the test fluid. Fluid flow was recorded with a Nikon D5300 ultra-high resolution single lens reflective (SLR) camera at 60 fps from a top-down view (Nikon, Tokyo, Japan).

Videos of flow dynamics were analyzed as described by Tokihiro et al.^33^ In brief, a Python program converted the videos into image sequences, with still images pulled every 10-200 frames depending on the length of the video. The segmented line and measure tools in ImageJ (National Institutes of Health) were used to quantify fluid displacement along the axial direction of a microchannel. The resulting measurements were exported as a CSV file.

## RESULTS AND DISCUSSION

### Design and fabrication of microchannels

Flow devices were designed to have a shallow rectangular open channel with rounded bottom corners to avoid capillary filament formation (Figure 1A).^34^ Due to the size and geometric constraints of the flow device, separate blocks were designed to be compatible with surface characterization instrumentation while preserving important characteristics of the channel. ‘Profilometry blocks’ with channels of the same height and width were designed for SEM and stylus profilometry. Profilometry blocks featured two full channels for surface texture analysis of the channel floors flanked by two half channels for analysis of the channel walls (Figure 1B). ‘Contact angle blocks’ were designed to create large, geometrically unconstrained surfaces for goniometry. The contact angle blocks featured a surface that mimics the channel floor and perpendicular surfaces to mimic the channel walls (Figure 1C).

All designed parts were fabricated with a laser SLA 3D printer. The parts were oriented with channels perpendicular to the laser rastering path (Figure 1D). For contact angle blocks, the long side of the ‘floor’ was set perpendicular to the laser path. To study the effect of print orientation on channel surface texture and capillary-driven fluid flow, the parts were angled with the length of the channel tilted at 0°, 15°, 45°, and 90° (Figure 1E). All parts were printed with the channel facing away from the build platform and supporting material with a z-layer height of 0.1 mm. A smaller layer height would increase the resolution of the channels, but 0.1 mm was selected because it is most feasible, efficient, and accessible for microfluidic fabrication users by reducing print time and extending the life of the 3D printer.

### Comparison of channel topography between print angles

To qualitatively compare the surface texture between channels printed at different angles, we used SEM to image the channel surface of profilometry blocks printed at 0°, 15°, 45°, and 90°. Representative SEM images show remarkable qualitative differences in the surface texture for channel floors. For channels printed at 0°, the channel floor has minimized surface texture with randomly distributed micro-bumps whereas channels printed at 15°, 45°, and 90° degrees have homogeneous, repetitive, and evenly distributed reliefs that run perpendicular to the channel (Fig. 2A-D). The relief pattern forms as a result of channel formation through successive *z*-layers.

**Figure 2.**
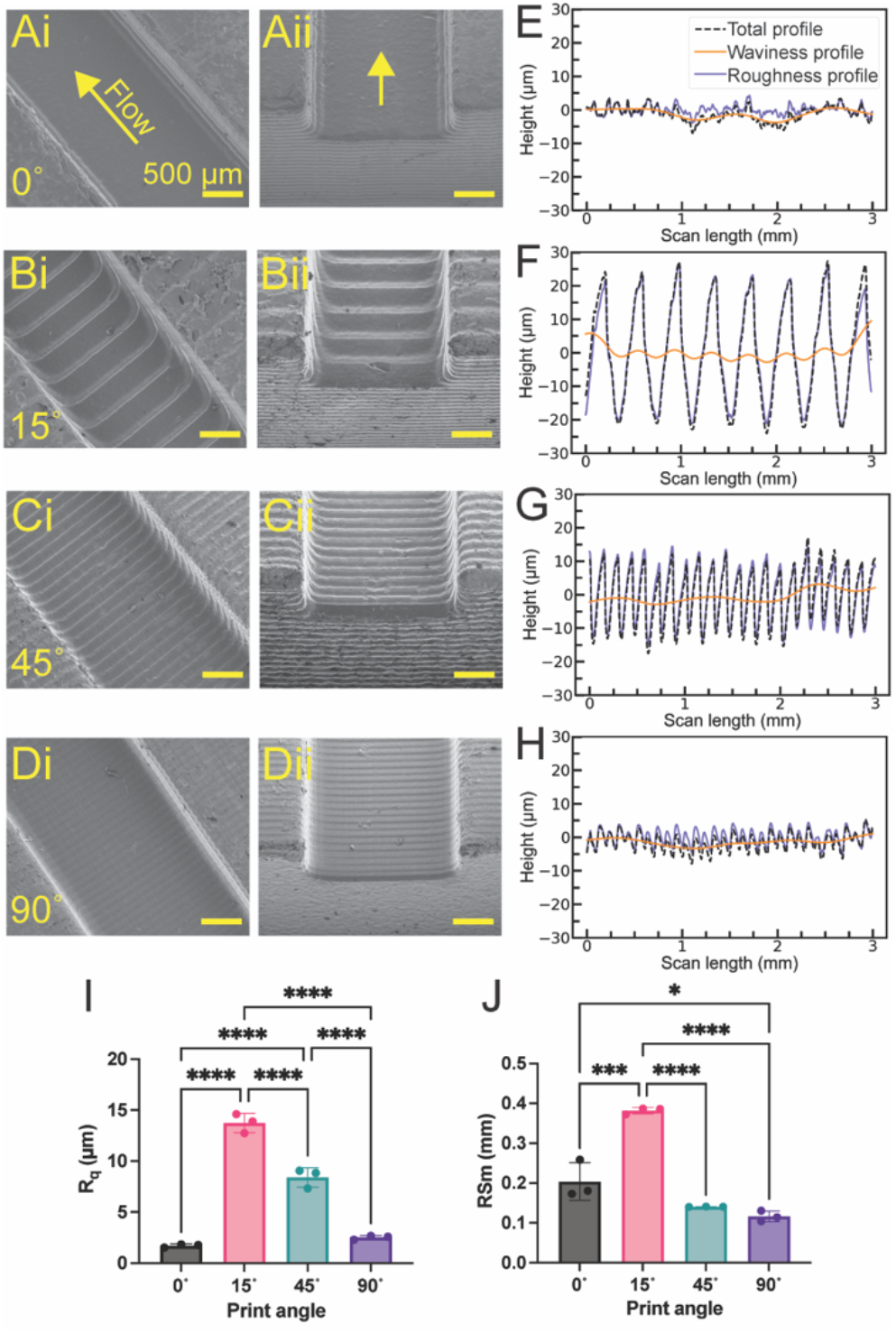
Print orientation greatly impacts surface texture and roughness of channel floors. (A-D) SEM images of profilometry block channels printed at 0° (A), 15° (B), 45° (C), and 90° (D) highlighting the channel floor (i) or the channel shape from an isometric view (ii). The arrow indicates the direction of flow. (E-H) Representative plots of channel topography measured along the direction of flow for channels printed at 0° (E), 15° (F), 45° (G), and 90° (H). The total profile is shown by the black dotted line, and the waviness and roughness profiles are shown in the orange solid line and slate blue solid line, respectively. (I and J) The differences between the topography plots have been quantified with R_q_ (RMS roughness) (I) and RSm (peak mean spacing) (J). Each point represents the average of 4 measurements on one independently printed part (n=3), and the error bars represent the SD. Statistical analysis was performed using a one-way ANOVA with Tukey’s multiple comparisons post hoc test. *p ≤ 0.05, ***p ≤ 0.001, ****p ≤ 0.0001.

To quantify the change in surface texture, the *z*-profile of the profilometry floors was measured in the direction of flow with a stylus profilometer. Representative profilometry scans are shown in Figures 2E-H, with the *z*-profile represented by the total profile. A Gaussian Regression filter was applied with a long cutoff of 0.8 mm to filter out low spatial frequencies associated with millimeter scale surface deviations (waviness) and select for high spatial frequencies associated with micrometer scale deviations (roughness). Spatial frequencies that pass the long cutoff filter generate the waviness profile, and spatial frequencies that are rejected generate the roughness profile. With the roughness profiles for each profilometry scan, we used two parameters to quantify differences in topography between various print angles - root-mean-square roughness (R_q_) and mean spacing between roughness profile irregularities (RSm). R_q_ captures the height deviations of peaks and valleys from the mean line, whereas RSm describes the frequency of peak and valley occurrence. 15° printed channel floors had the highest R_q_ and RSm, demonstrating that peak deviations were large but occurred less frequently (Fig. 2I and J). In contrast, 45° printed channel floors had the second highest R_q_ and a RSm value of less than half of the 15° printed channel floor (Fig. 2I and J). Therefore, the peak deviations for the 45° printed channel floors were smaller but occurred more than twice as frequently when compared to the 15° printed channel.

To capture the differences between the various print angles with a parameter that considers both peak deviation and frequency, we calculated the tortuosity of each profilometry scan. Tortuosity is defined as

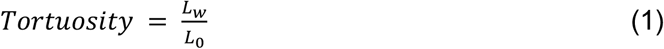

where L_w_ is the real length along the surface and L_0_ is the straight-line, theoretical length. The values for tortuosity are shown in Table 1, with high tortuosity indicating a longer real channel length. The tortuosity of the 45° printed channel floors was the greatest, implying that fluids would have a longer effective travel distance in a channel printed at 45° as opposed to a channel printed at 0°.

**Table 1.** Tortuosity (τ) of microchannel floors and walls as calculated by Eq. 1. Values are expressed as mean ± standard deviation (SD).

| Print angle | 0° | 15° | 45° | 90° |
| --- | --- | --- | --- | --- |
| Floor | 1.40 ± 0.08<sup>a</sup> | 2.63 ± 0.14 | 3.69 ± 0.38 | 1.69 ± 0.04<sup>a</sup> |
| Wall | 1.52 ± 0.01<sup>b</sup> | 1.55 ± 0.05<sup>b</sup> | 1.64 ± 0.08<sup>b</sup> | 1.62 ± 0.08<sup>b</sup> |

<sup>a,b</sup>Note: No statistical difference was found between values with like notations. There were statistical differences found between the remaining values (Fig. SI.4.1). Statistical analysis was performed using a one-way ANOVA with Tukey's multiple comparisons post hoc test.

Similar analysis of surface texture and roughness was conducted for the channel walls of profilometry blocks printed at 0°, 15°, 45°, and 90°. The channel walls were also affected by print angle as seen in representative SEM micrographs (Fig 3A-D). While the surface texture of the channel walls looked qualitatively different at varying print angles, the difference between R_q_ values was smaller in magnitude than the difference between R_q_ values of the channel floors (Fig 3I). The tortuosity values of the channel walls were also similar regardless of print angle (Table 1). Despite the qualitative differences observed between print angles, stylus profilometry measurements did not capture any quantitative differences. A possible explanation is that the 2D stylus profilometer only scans over a length in a single direction. A different surface measurement method that analyzes surface topography over a given area in multiple directions may reveal more quantitative differences. This could be explored in future work with a 3D profilometer. Overall, in our study, print angle has a greater effect on the texture of the floors than the walls. Altogether, the data show that print orientation can greatly alter the surface topography of the channel floor and moderately alter the topography of the channel walls.

**Figure 3.**
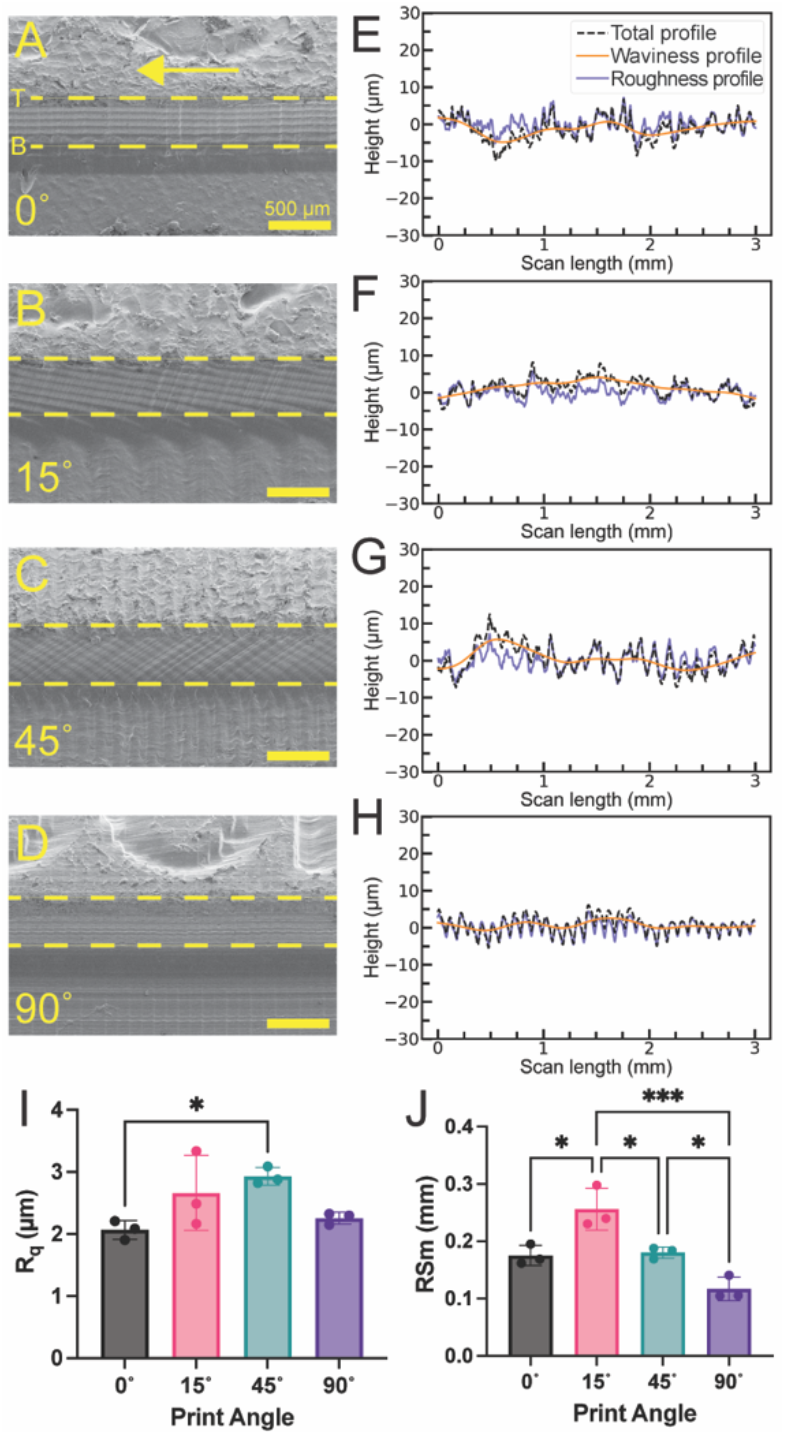
Print orientation moderately impacts surface texture and roughness of channel walls. (A-D) SEM images of profilometry block channels printed at 0° (A), 15° (B), 45° (C), and 90°. The arrow indicates the direction of flow. The dashed lines indicate the top (‘T’) and bottom (‘B’) of the channel walls. (E-H) Representative plots of channel topography measured along the direction of flow for channels printed at 0° (E), 15° (F), 45° (G), and 90° (H). The total profile is shown by the black dotted line, and the waviness and roughness profiles are shown in the orange solid line and slate blue solid line, respectively. (I and J) The differences between the topography plots have been quantified with R_q_ (I) and RSm (J). Each point represents the average of 4 measurements on one independently printed part (n=3), and the error bars represent the SD. Statistical analysis was performed using a one-way ANOVA with Tukey’s multiple comparisons post hoc test. *p ≤ 0.05 and ***p ≤ 0.001.

### Comparison of wetting behavior between print angles

It has been well established that small-scale surface roughness accentuates the wetting characteristics of a solid substrate.^35,36^ According to Wenzel’s law, very-small-size roughness causes wetting of an aqueous solution to increase for a hydrophilic surface and decrease for a hydrophobic surface. However, in the presence of large-size roughness (presented in the form of grooves and reliefs) the droplet footprint is not circular, and Wenzel’s law does not apply. The droplet adopts a non-circular shape that minimizes its surface energy, guided by the relief.^37,38^ Fluid wetting behavior can be quantified by measuring the contact angle, which is the angle between a line tangent to the air-liquid interface and the solid surface. When the measurement is taken on an ideal (rigid, smooth, chemically homogeneous, and nonreactive) surface, the static contact angle is measured. For 3D-printed channels, the surface is nonideal with the presence of peaks and valleys created by roughness reliefs; the apparent contact angle between the line tangent to the air-liquid interface and a line representing the nominal solid surface is measured instead.^39^ The contact angle is an important parameter that determines the potential for spontaneous capillary flow (SCF). The condition for SCF in a monolithic channel is

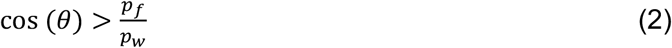

where the contact angle, θ, is the same for every solid surface. *p*_*f*_ and *p*_*w*_describe the air-liquid interfaces (free perimeter) and solid-liquid interfaces (wetted perimeter) of the channel cross section, respectively.^27,29^

To assess the effect of print orientation-mediated surface texture and roughness on droplet spreading behavior, we used a drop shape analyzer (goniometer) to measure the apparent contact angle of a droplet of glycerol solution on plasma-treated contact-angle blocks printed at 0°, 15°, 45°, and 90°. The properties of the glycerol solution are described in Table 2. Apparent contact angle measurements were taken on both the ‘floor’ and ‘wall’ sides of the contact angle block (Fig 4A). Measurements on each side were taken from two viewpoints: camera angle 1 captures the contact angle as the droplet spreads in the direction parallel to flow and camera angle 2 captures spreading perpendicular to flow (Fig 4B and C).

**Table 2.** Experimental solution properties. Values are expressed as mean ± SD.

| Solution | Concentration | Surface tension (mN/m) | Viscosity (mPa·s) | Contact angle on smooth SLA resin (°) |
| --- | --- | --- | --- | --- |
| Glycerol | 0.644 (w/w) | 70.6 ± 3.8 | 15.7 ± 0.7 | 37.3 ± 4.0 |
| Agarose | 1.5% (w/v) | 70.4 ± 2.5 | 10.9-12.3<sup>a</sup> | 29.7 ± 8.6 |
| Collagen | 3 mg/mL | 67.7 ± 1.4 | 17.2-18.3<sup>a</sup> | 24.4 ± 1.2 |

<sup>a</sup>Note: Viscosity is dependent on shear rate (Fig. SI.4.3). Viscosity values listed are based on the expected shear rate in open channels (in the range 10-50 s<sup>-1</sup>, corresponding to velocities in the range 1 to 5 mm/s).

**Figure 4.**
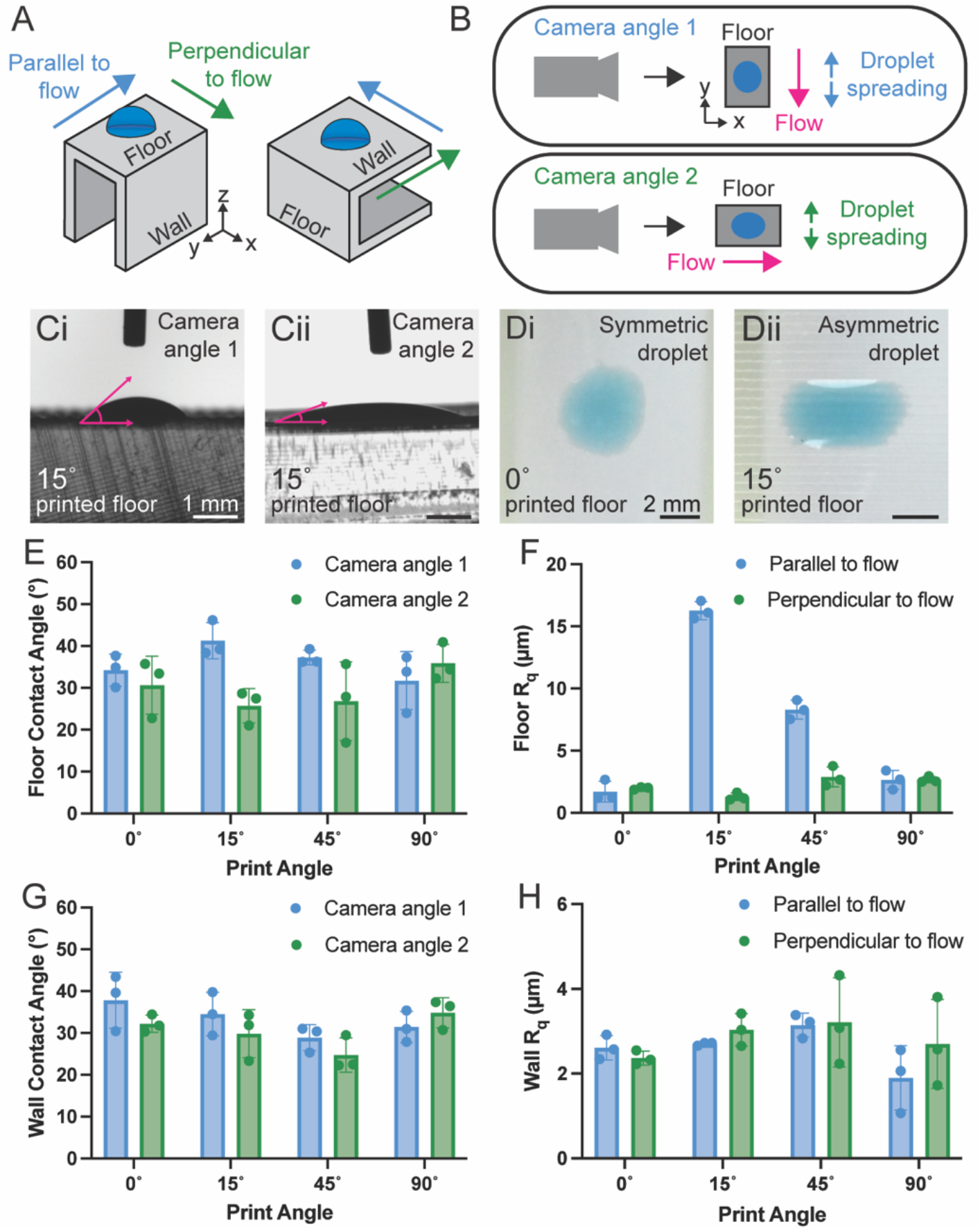
The resulting surface texture of varying print orientation can cause wetting asymmetry. (A) Schematic depicting contact angle block orientation to measure droplet spreading on the channel floor or wall. Due to specific orientation during printing, the long side of the block floor corresponds to the direction parallel to flow (blue arrows) and the short side of the block floor corresponds to the direction perpendicular to flow (green arrows). (B) Schematic illustrating block orientation during droplet shape analysis. In camera angle 1, the analyzer camera measures the contact angle as the droplet spreads parallel to flow. In camera angle 2, the analyzer camera measures the contact angle as the droplet spreads perpendicular to flow. (C) Side view images from a drop shape analyzer (goniometer) of glycerol solution droplets on a 15° printed block floor from camera angle 1 (i) and camera angle 2 (ii). (D) Top-down images of glycerol solution droplets on a 0° (i) and 15° (ii) printed block floor. (E and G) The contact angle of glycerol solution on the channel floor (E) and wall (G) was measured from both camera directions. (F and H) The R_q_ of the channel floor (F) and wall (H) were measured both parallel and perpendicular to flow. (E-H) Each point represents the average of 3 measurements on one independently printed part (n=3), and the error bars represent the SD.

We found that droplets formed a circular footprint on contact angle block floors printed at 0° and 90° and a non-circular footprint for 15° and 45° (Fig 4D and E). When the droplet footprint is non-circular, apparent contact angle measurements taken from camera angles 1 and 2 will differ greatly; wetting is asymmetric, and fluid spreading is greater in the direction where the apparent contact angle is smaller. To investigate the relationship between wetting behavior and surface roughness, stylus profilometry was conducted for the contact angle block floors with the stylus moving parallel to flow and perpendicular to flow, which revealed symmetric roughness for 0° and 90° printed floors printed and asymmetric roughness for 15° and 45° printed floors (Fig 4F). Increased roughness in the direction parallel to flow did not correlate with a smaller contact angle for camera angle 1 on the 15° and 45° floors. Due to the large size of the roughness reliefs compared to the droplet size (Fig. SI.4.2), Wenzel’s law does not apply; instead, the reliefs act as microchannels (within the larger microchannel) for the fluid to wick along. In the context of the flow devices, these reliefs span perpendicular to the direction flow; thus, presenting the opportunity for fluid pinning along the length of the relief (Fig 4Dii). Fluid pinning would impede flow along the length of the channel and decrease flow velocity. This could be advantageous if attenuated fluid velocity is needed for applications such as delay time for reaction mixing.

Contact angle block walls were found to have more symmetrical droplets and roughness regardless of print angle (Fig 4G and H). There were variations in the measured apparent contact angles across the different print angles, but the association between apparent contact angle and R_q_ across the different print angles was not clear. As discussed before, a bi-directional measurement of surface topography could better capture the qualitative differences observed in the texture of the channel walls, and this is the subject of future work. A parameter associated with bi-directional measurement could explain the observed variation in apparent contact angle with respect to print angle. The results shown here support the finding that print angle has a larger impact on the characteristics of the channel floor compared to the channel wall. Further, print orientation-mediated surface texture affects wetting behavior, a parameter related to capillary-driven flow (Eq. 2).

### Comparison of capillary driven flow between print angles

Open microfluidic channels have been commonly leveraged to pattern cell-embedded hydrogel precursors and create more physiologically relevant 3D cell culture models.^20–26^ To better understand how interactions with a textured channel surface may affect capillary-driven flow of biologically relevant complex fluids, flow tests were conducted with glycerol solution, agarose precursor solution, and collagen precursor solution in flow devices (Fig. 1A) printed at 0°, 15°, 45°, and 90°. Glycerol solution was chosen due to its aqueous nature and was formulated to have a similar viscosity to the agarose and collagen precursor solutions (Table 2, Fig. SI.4.3). Agarose and collagen precursors were selected for their common use in microfluidic hydrogel patterning applications.^24,27,40,41^ The properties of all test solutions are listed in Table 2.

To ensure that the channel height and width remained similar regardless of print angle, unique channel dimensions were designed for each print angle so that all channel dimensions would be within 5% error (Fig. SI.4.4, Table SI.3.1). Since rougher surfaces produce more friction, we expected to observe the slowest flow in 15° printed channels, which have the highest floor R_q_. Contrary to expectations, flow of glycerol solution was much slower in flow devices printed at 45° compared to 0°, 15°, and 90° (Fig 5A and B, Fig. SI.4.5). Rather than R_q_, which only considers peak height deviation, we found that floor tortuosity (Table 2), which considers both peak height deviation and frequency, acted as a better predictor of slow flow.

**Figure 5.**
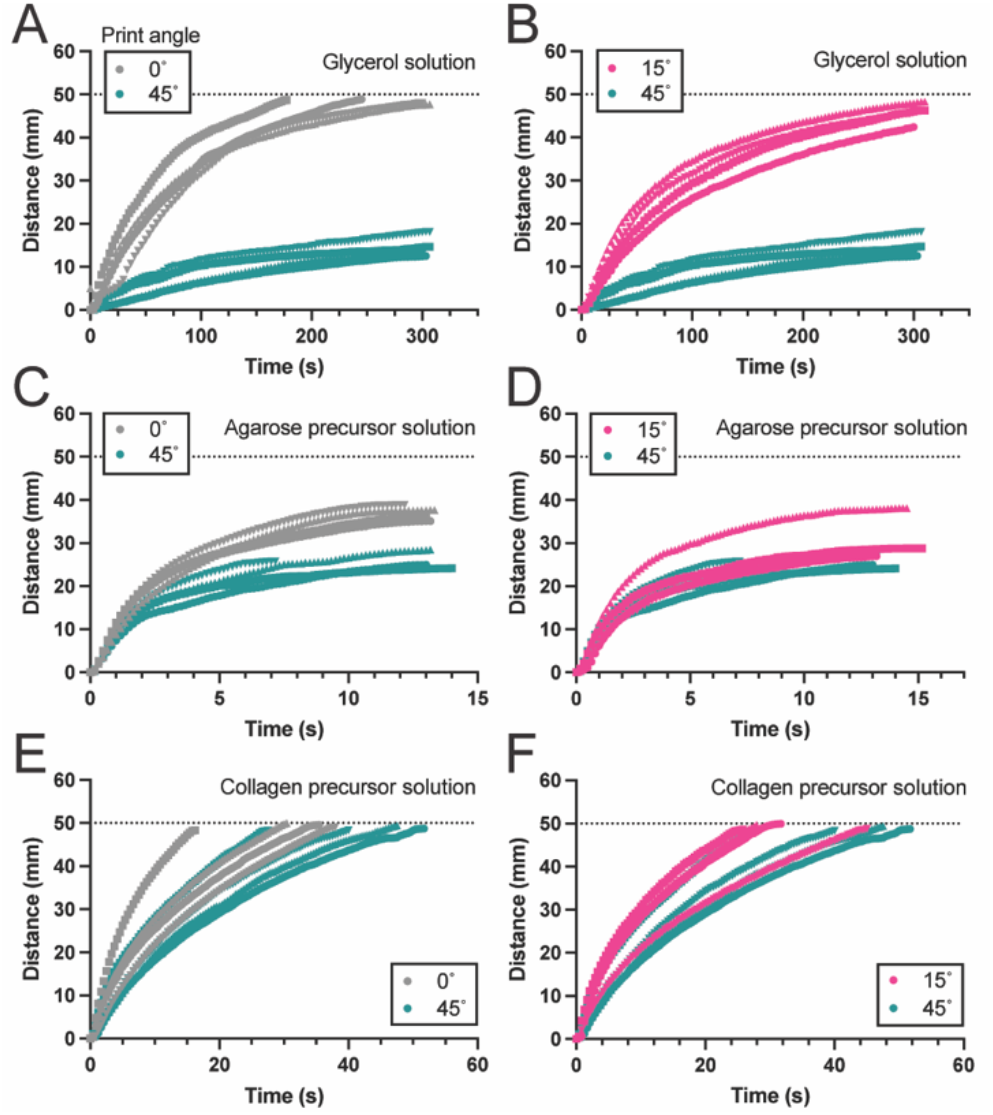
Print orientation greatly affects SCF of glycerol solution and moderately affects flow of agarose and collagen precursor solutions. Flow of glycerol solution is compared between 0° and 45° microchannels (A) and 15° and 45° microchannels (B). Flow of agarose precursor solution is compared between 0° and 45° microchannels (C) and 15° and 45° microchannels (D). Flow of collagen precursor solution is compared between 0° and 45° microchannels (E) and 15° and 45° microchannels (F). The black dotted lines represent the end of the channel. Each colored line represents flow in an independently printed flow device (n=4).

Flow with agarose and collagen precursor solutions was much faster than flow with glycerol solution, and the effect of print angle on flow velocity was moderated (Fig 5C and D, Fig. SI.4.6, Fig. SI.4.7). However, for agarose precursor, a solution that solidifies rapidly at room temperature, print angle remained an important consideration as the overall travel distance of the fluid was about 30% less in channels printed at 45° compared to 0° (Fig 5C). The effects of print angle are likely minimized for agarose and collagen precursors due to the lower contact angle of the solutions on plasma-treated 3D printed resin (Table 2), as a low contact angle facilitates faster flow. Additionally, the surface tension and viscosity, other parameters that affect flow, are similar between the three solutions. While the measured contact angle of agarose precursor solution is higher than collagen precursor solution, this measurement is likely inflated due to the rapid gelling of the agarose precursor. When still images of the flow videos were compared, the first drop of test solution was flatter, or more wetted, for agarose precursor compared to collagen precursor (Fig. SI.4.8). This would explain why faster flow was observed for agarose precursor solution compared to collagen precursor solution.

Taken as a whole, our results show that print angle can greatly affect capillary driven flow depending on the contact angle of the fluid. Moreover, R_q_ is not a reliable predictor of the effect of surface topography on flow; instead, floor surface tortuosity should be considered. These results have important implications for researchers designing microfluidic-based hydrogel patterning platforms. For platforms designed to pattern quick gelling hydrogel precursors such as fibrin, a print orientation that minimizes floor texture, such as 0° print angle, may be recommended to ensure the hydrogel precursor fills the entire channel before gelling. On the other hand, for some platforms that require precisely timed and controlled patterning, a print orientation that increases floor texture, such as 45° print angle, and causes delayed fluid velocity may be preferred.

### New approaches for approximating effect of surface roughness

Current theoretical models for capillary-driven flow dynamics cannot accurately predict the effects of surface texture and roughness on flow in open channels. Additionally, R_q_ and R_a_ (average roughness) measurements are conventionally used to report the surface qualities of various microchannel fabrication techniques,^14,16,31,32^ but these measurements were not sufficient to predict roughness-mediated flow patterns. Thus, work to develop new theoretical models that account for surface roughness effects beyond R_q_ and R_a_ is needed to bridge our understanding on how roughness phenomena affects flow.

The travel distance (*z*) as a function of time (*t*) in a smooth, open microchannel is modeled by the modified Lucas-Washburn-Rideal (mLWR) law,

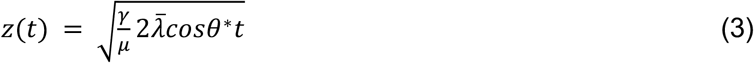

where #and #are the surface tension and viscosity, respectively; 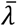 is the friction length, which describes the effect of wall friction; and θ* is the generalized Cassie angle, which accounts for the effect of the free surface.^42,33^ In Figure 6A, the mLWR model for collagen precursor solution (orange line) is compared with an experimental replicate of collagen precursor solution flowing in a 90° printed flow device (purple triangles). While the experimental data show slower flow than predicted by the model, the flow motion in laser SLA-based channels (where roughness is relatively small, homogeneous, and repetitive along the channel length) still approximately follows a square-root law 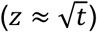. Flow occurs more slowly in 3D-printed channels than predicted by the mLWR law, consistent with previous findings.^32^ A few explanations have been proposed, all related to the increased surface roughness typical of 3D printing: (1) transient, partial pinning of the meniscus on surface “micro-bumps”^32,43–45^ (Fig. 6B) and (2) enhanced wall friction caused by the interaction with rough surfaces.^46,47^ Further work is still needed to adapt the mLWR law to account for roughness-induced effects. Below, we propose that altered flow dynamics due to roughness is governed by two parameters: the apparent average friction length and the average apparent contact angle.

**Figure 6.**
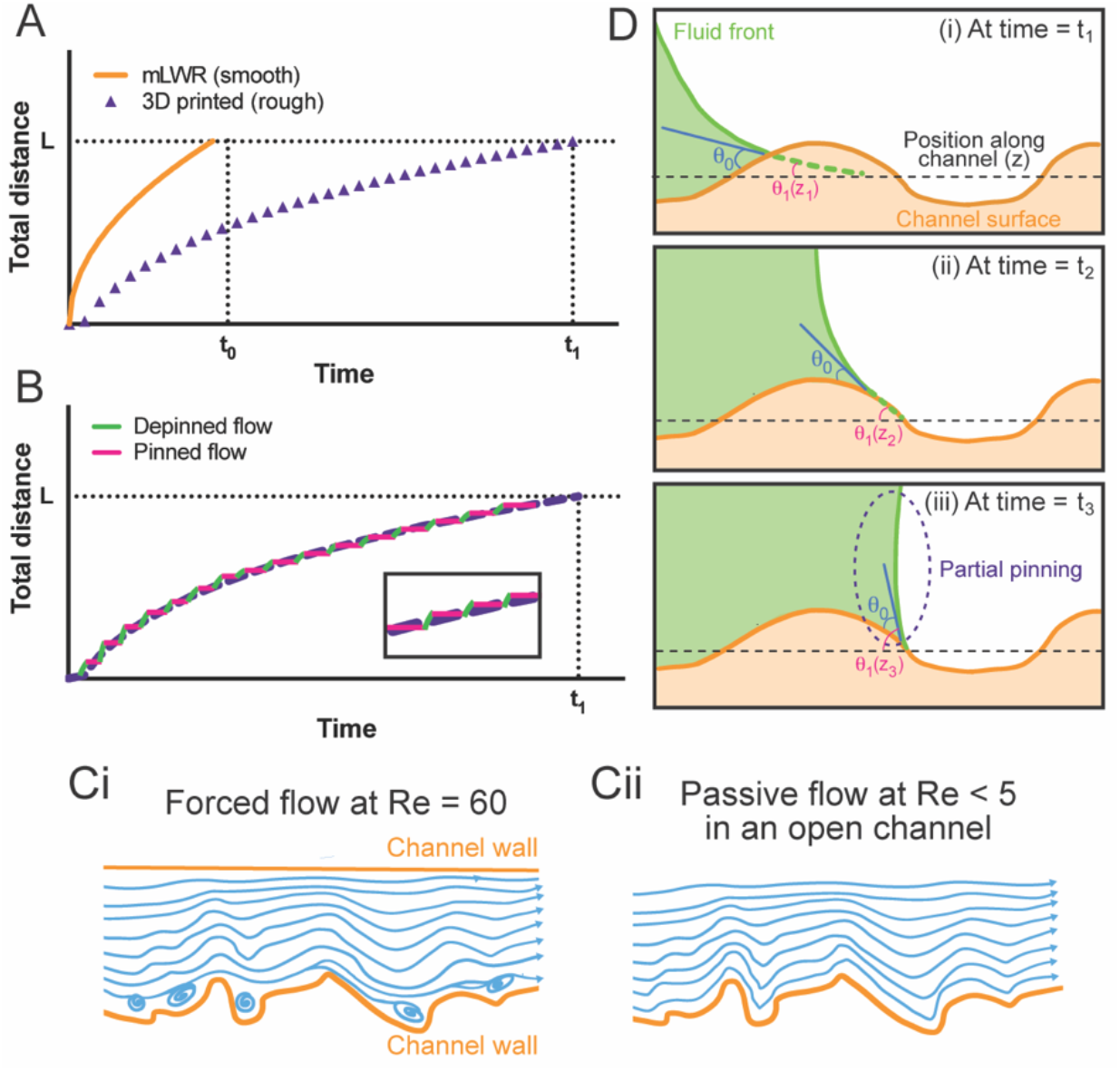
Capillary flow on homogeneously rough surfaces. (A) Plot comparing the mLWR model for flow of collagen precursor solution in a smooth channel (orange line) compared to one experimental replicate of flow of collagen precursor solution in a 90°printed channel (purple triangles). The black dotted horizontal line, L, represents the end of the channel. (B) Theoretical model of travel distance vs. time in the case of small-scale roughness and temporary pinning. The green and pink lines represent the fluid travel in the depinned and pinned state, respectively. Inset shows magnified view. C) Schematic of a laminar flow (i) with recirculation (Re = 60 in closed channel with pumping) and (ii) without recirculation (Re < 5 in open channel without pumping). Blue lines depict fluid streamlines. (D) Schematic of the static contact angle (θ_#_, blue) and apparent contact angle (θ_#_(z), pink) as the fluid front moves along a homogenous rough surface when time is t_1_ (i), t_2_ (ii), and t_3_ (iii).

As stated previously, the average friction length (#) describes the effect of wall friction. For laminar forced flows in closed microchannels at Reynolds number (Re) between 10 and 100, the effect of roughness on wall friction is noticeable and linked to the size of the reliefs.^47^ An argument for the increased wall friction was that the reliefs act as constrictions on the flow, leading to fluid recirculation (Figure 6Ci).^47^ In the case of capillary flow in open channels, pinching is weak due to the open surface and the Reynolds number is still lower–in the range 1 to less than 10. At these low Reynolds numbers, occurrence of recirculation disappears, and the streamlines closely follow the wall (Figure 6Cii). Hence, wall friction increases due to the longer contact length between the fluid and the wall. The increase in total wall length can be described by tortuosity, #(Eq. 1). We propose correcting #with #to obtain the apparent average friction length for channels with homogeneous, repetitive, and symmetrical roughness where

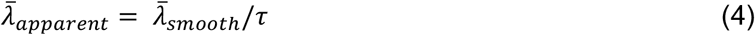

A tentative explanation can be found in SI.1. As tortuosity increases, apparent average friction length decreases; and distance traveled *z(t)*, as predicted by the mLWR law in Equation 3, also decreases. This model would explain why flow velocity was slower in the 45° printed channels compared to the 15° printed channels. Despite the larger reliefs and greater R_q_ present in 15° printed channels, 45° printed channels had tighter spacing between the reliefs and greater surface tortuosity.

The apparent contact angle is linked to the shape of the advancing meniscus at the front of the flow, locally changing due to the roughness reliefs on the walls. Previous work has documented slower fluid velocity in 3D printed channels due to the occurrence of successive temporary local pinning of capillary flow (Fig 6B).^32,43–45^ In a rough channel with continuous local pinning and depinning, the apparent contact angle of the fluid front, θ_#_, changes depending on where the meniscus is located in relation to the roughness reliefs (Fig 6D). In the mLWR law, the effect of contact angle is represented by the Cassie angle, #θ^#^. The Cassie angle is only a function of the channel cross section geometry and the static contact angle (θ_#_). To better model capillary flow in a channel with homogeneous roughness, we propose considering the average of apparent contact angles along the flow path in the axial direction (Fig 6D). We speculate that the average apparent contact angle can be mathematically determined from the ‘rough’ mLWR law and the apparent average friction length. A full derivation for open rectangular channels can be found in SI.2.

In future work, we will validate these proposed theoretical variables, apparent average friction length as corrected by #and average apparent contact angle, through experiments with well-characterized fluids for fundamental microfluidic studies (e.g., water, nonanol), comparison experiments with smooth 3D-printed channels, and computational modeling. We anticipate that the introduction of these parameters may improve theoretical models of capillary-driven flow in rough open channels. Nevertheless, our current work presents an advance in our understanding on the effects of small, homogeneous roughness inherent to laser SLA printing on surface-tension driven flow.

## CONCLUSIONS

This paper presents advances in two areas: (1) proposing new considerations for improving fundamental understanding of fluid flow in rough, open-capillary systems and (2) providing a practical framework to guide laser SLA-based microfluidic fabrication through systematic characterization of various print orientations. To briefly summarize the previous section, we propose the consideration of apparent average friction length, as modified by #, and average apparent contact angle to improve theoretical models for estimating capillary driven flow in homogeneously rough, 3D-printed open microchannels. Future work will further explore these parameters with conventional fluids used in fundamental microfluidic studies such as water and nonanol.

Concerning creating a practical framework, we systematically characterized microchannels printed at 4 different angles: 0°, 15°, 45°, and 90°. Our findings demonstrated that tilting the length of the channel has substantial effects on surface morphology, fluid wetting behavior, and capillary-driven flow that warrant consideration from microfluidic users. Channel floors printed at 15° had the greatest R_q_, whereas channel floors printed at 45° had the greatest tortuosity. Both print angles resulted in asymmetric roughness, which led to asymmetric wetting behavior where the fluid spreads further parallel to the roughness reliefs. Channel walls had qualitative texture differences captured by SEM that could not be quantified through 2D profilometry; future work will explore 3D topography measurement methods. Printing at 45° caused delayed channel filling for glycerol solution as well as agarose and collagen precursor solutions, and the effects of print orientation were more pronounced for fluids with higher contact angles and hydrogel precursors that gel quickly. Reduced fluid flow due to print angle was better predicted by #, rather than R_q_. These findings are summarized in Table 3.

**Table 3.** Summary of findings on comparison of print angle with evaluated parameters.

|  | 0° print angle | 15° print angle | 45° print angle | 90° print angle |
| --- | --- | --- | --- | --- |
| Channel texture (floor) | No visible banding<br>Lowest $R_q^a$<br>Lowest $\tau^b$ | Perpendicular banding<br>Highest $R_q$<br>Highest RSm | Perpendicular banding<br>Highest $\tau$ | Perpendicular banding<br>Lowest $R_q^a$<br>Lowest $\tau^b$ |
| Channel texture (wall) | Parallel layering | Cross hatch layering | Cross hatch layering | Parallel and perpendicular layering |
| Fluid wetting (floor) | Symmetric wetting | Asymmetric wetting | Asymmetric wetting | Symmetric wetting |
| Fluid wetting (wall) | Symmetric wetting | Symmetric wetting | Symmetric wetting | Symmetric wetting |
| Capillary driven flow | Fastest for agarose precursor |  | Slowest for all fluids |  |
| Recommendations | Recommended for fast-gelling hydrogel precursors |  | Recommended for applications that require delay time or controlled flow |  |

<sup>a,b</sup>Note: No statistical difference was found between values with like notations.

Our results provide a guide for microfluidic users to best fabricate their devices with laser SLA depending on their desired fluid velocity for applications with simple fluids or complex hydrogel precursors. For applications that require delay time or slower and more controlled flow, we suggest printing microchannels at a print angle that increases floor texture such as 45°. For flowing quick gelling hydrogel precursors, we suggest a print angle that decreases floor texture such as 0°. We also recommend that microfluidic users be consistent with their selected print orientation throughout the device development process. Our findings on the effects of laser SLA print orientation may be applicable to users of DLP-SLA and LCD-SLA due to the layer-by-layer nature of all three fabrication methods. Our work also presents tortuosity as an alternative method for quantifying surface texture, a better predictor of fluid velocity, and a potential consideration to create better theoretical models for capillary flow in rough channels–all of which is germane to any fabrication method.

## ACKNOWLEDGEMENTS

Research reported in this publication was supported by the National Institutes of Health (NIH) National Institute of General Medical Sciences Grant R35GM128648 (A.B.T.), the Fulbright Program (L.A.M.), the University of Washington Department of Chemistry Summer Research Award and Levinson Emerging Scholars Award (D.W.H.C.), and the National Center for Advancing Translational Sciences Grant TL1TR002318 (J.C.T. and L.G.B.). Part of this work was conducted at the Molecular Analysis Facility, a National Nanotechnology Coordinated Infrastructure (NNCI) site at the University of Washington, which is supported in part by funds from the National Science foundation (awards NNCI-2025489, NNCI-1542101), the Molecular Engineering & Sciences Institute, and the Clean Energy Institute. We also acknowledge the M.J. Murdock Diagnostics Foundry for Translational Research. The content is solely the responsibility of the authors and does not necessarily represent the official views of the National Institutes of Health or other funding bodies.

