## Supporting Information for "From Fabrication to Flow: Impact of Print Orientation on Surface Qualities and Capillary-Driven Flow in Laser SLA-based Open Microchannels"

### Table of Contents

| Section | | Page |
|---|---|---|
| SI.1 | Model for wall friction based on line tortuosity along a surface | S2 |
| SI.2 | Model for the apparent contagct angle along a rough surface | S5 |
| SI.3 | Engineering drawings and channel cross section dimensions | S8 |
| SI.4 | Additional experimental data | S11 |
| SI.5 | Supporting information references | S18 |

**Additional supporting information included:**

Video: flow experiment with glycerol solution in 0˚ printed device (.mp4)
Video: flow experiment with agarose precursor solution 0˚ printed device (.mp4)
Video: flow experiment with collagen precursor solution 0˚ printed device (.mp4)
Design file: 0˚ print angle flow device (.STL)
Design file: 15˚ print angle flow device (.STL)
Design file: 45˚ print angle flow device (.STL)
Design file: 90˚ print angle flow device (.STL)
Design file: profilometry block (.STL)
Design file: contact angle block (.STL)

### SI.1. Model for wall friction based on line tortuosity along a surface

The notations used in this section are listed in Table SI.1.1.

Table SI.1.1. Notations

| Name | Notation | Unit | Remarks |
|---|---|---|---|
| Tortuosity | $\tau$ | unitless | |
| Length | $L$ | mm | Length along the axial direction of the channel |
| Real length | $L_w$ | mm | Real length along the surface |
| Straight-line length | $L_0$ | mm | Straight-line length along the surface |
| Perimeter (wetted) | $p_w$ | mm | Perimeter of channel cross section with liquid-solid interface |
| Perimeter (free) | $p_F$ | mm | Perimeter of channel cross section with liquid-air interface |
| Travel distance | $z$ | mm | Middle of advancing meniscus in smooth channel |
| Travel distance (rough) | $z^*$ | mm | Middle of advancing meniscus in rough channel |
| Surface tension | $\gamma$ | mN/m | Property of test fluid |
| Viscosity | $\mu$ | mPa·s | Property of test fluid |
| Average friction length | $\bar{\lambda}$ | mm | |
| Apparent average friction length | $\bar{\lambda}^*$ | mm | |
| Static contact angle | $\theta$ | rad | Static contact angle |
| Generalized Cassie angle | $\theta^*$ | rad | Generalized Cassie angle for open rectangular channel |
| Time | $t$ | s | |
| Pressure | $P$ | mPa | |
| Velocity | $V$ | mm/s | Velocity in smooth channel |
| Velocity (rough) | $V^*$ | mm/s | Velocity in rough channel |
| Cross-sectional area | $S$ | $mm^2$ | Cross-sectional area of channel |

Laminar wall-friction at the microscale has mainly been investigated for forced flows. These studies showed that Moody's diagram requires adjustments for roughness effects when the Reynolds number is below 100.[1–4] Both the relative roughness—defined as the ratio of the characteristic roughness height to the channel size—and the channel's aspect ratio were found to influence the pressure drop. For example, Jia et al.[4] demonstrated that micro-scale recirculating flow structures can develop in forced flows with Reynolds numbers between 15 and 100 (Fig. SI.1.1A). These recirculation zones increase viscous dissipation, thereby leading to a higher-pressure drop. In the case of capillary flow, especially in open channels, the Reynolds number is still lower, in the range 1 to 10.[5] At these low Reynolds numbers, occurrence of recirculation disappears, and the streamlines follow closely the wall (Fig. SI.1.1B). Hence, wall friction increases due to the longer contact length between the fluid and the wall. This length is used to define the axial wall tortuosity by extension of the work of Kozeny[6] and Carman[7] in the years 1927 to 1937 for porous media where

$$\tau = L_w/L_0, \qquad \text{(SI1.1)}$$

$L_w$ being the real length along the surface while $L_0$ is the straight-line length.

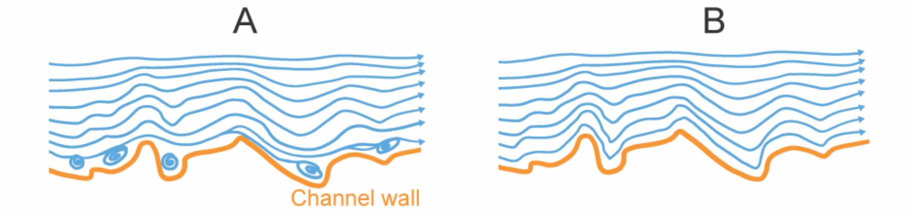

**Figure SI.1.1**. (A) Wall recirculation in a forced flow at Reynolds 60. (B) Sketch of wall streamlines at very low Reynolds numbers.

Thus, the present model for wall friction in an open channel is based on the hypothesis that the wall contact surface is only increased by the wall roughness and the average friction length being the same as that of the equivalent smooth channel. Let us be reminded that, in an open channel, an average friction length has been defined to account for the wall friction, so that the modified Lucas Washburn law is

$$z = \sqrt{\frac{\gamma}{\mu} 2\bar{\lambda} \cos\theta^* \ t}, \qquad \text{(SI1.2)}$$

where $\theta^*$ is the generalized Cassie angle and $\bar{\lambda}$ the average friction length (on the contour of the cross-perimeter). The length $\bar{\lambda}$ is only associated to the channel cross-sectional geometry. The generalized Cassie angle is

$$\cos\theta^* = \frac{p_w \cos\theta - p_F}{p_w + p_F}, \qquad \text{(SI1.3)}$$

where $p_w$ and $p_F$ are the wetted and free perimeter, respectively. The hypothesis that the average friction length stays identical is based on the observation that, if the roughness reliefs are small compared to the cross dimensions of the channel, the velocity profile is only slightly modified at the walls as shown in Figure SI.1.2.

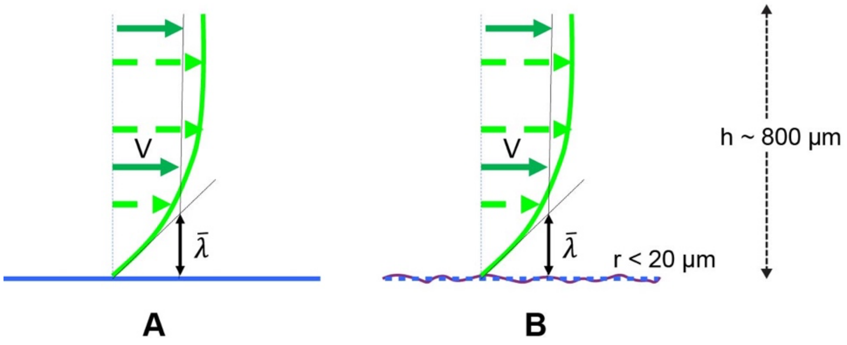

**Figure SI.1.2**. (A) Sketch of the flow profile (green lines) along a smooth wall (blue line). (B) In presence of roughness (purple solid line), the flow profile stays identical in the whole core flow and is only slightly modified close to the wall; hence, the hypothesis that $\bar{\lambda}$ remains approximately the same.

Based on these considerations, let us write the expression of the wall friction (pressure drop along the distance *L*)

$$\Delta P = \frac{p_w}{S} \int_0^{L\tau} \mu \frac{\partial v}{dy}\, dz \;, \tag{SI1.3}$$

where *S* is the cross-sectional area. Assuming the same average shear rate $\frac{\partial v}{dy} = \frac{V}{\bar{\lambda}}$, the pressure drop becomes

$$\Delta P = \frac{L_0 \tau\, p}{S}\, \mu \frac{V}{\bar{\lambda}} = \frac{L_0\, p}{S}\, \mu \frac{V}{\frac{\bar{\lambda}}{\tau}} \;. \tag{SI1.4}$$

It is deduced that the effect of roughness on friction is taken into account by using the apparent friction length

$$\bar{\lambda}^* = \bar{\lambda}/\tau \;. \tag{SI1.5}$$

The tortuosity $\tau$ being always larger than 1, the friction length is reduced by roughness and the flow is slower

$$z^* = \sqrt{\frac{\gamma}{\mu} 2 \frac{\bar{\lambda}}{\tau} cos\, \theta^*\;t} \;, \tag{SI1.6}$$

then the "rough" travel distance is related to the "smooth" travel distance by

$$z^* = \frac{1}{\sqrt{\tau}}\, z \;. \tag{SI1.7}$$

Consequently, the "roughness-corrected" velocity, $V^*$, is

$$V^* = \frac{1}{\sqrt{\tau}}\, V \;. \tag{SI1.8}$$

### SI.2. Model for the apparent contact angle along a rough surface

The notations used in this section are listed in Table SI.2.1.

Table SI.2.1. Notations

| Name | Notation | Unit | Remarks |
|---|---|---|---|
| Width | $w$ | mm | Width of channel |
| Height | $h$ | mm | Height of channel |
| Length | $L$ | mm | Length along the axial direction of the channel |
| Length | $L_1$ | mm | Length along the axial direction of the channel at $t_1$ |
| Static contact angle | $\theta_0$ | rad | |
| Apparent contact angle | $\theta_1$ | rad | |
| Average apparent contact angle | $\theta_{1,app}$ | rad | |
| Generalized Cassie angle | $\theta_1^*$ | rad | Apparent generalized Cassie angle for open rectangular channel |
| Travel distance | $z$ | mm | Middle of advancing meniscus |
| Surface tension | $\gamma$ | mN/m | Property of test fluid |
| Viscosity | $\mu$ | mPa·s | Property of test fluid |
| Average friction length | $\bar{\lambda}$ | mm | |
| Time | $t$ | s | |
| Time | $t_1$ | s | Arbitrary timepoint before fluid reaches the end of the channel |

If the contact angle between the liquid and the wall stays always the same, the presence of roughness induces the notion of “apparent contact angle”, as shown in Figure SI.2.1.

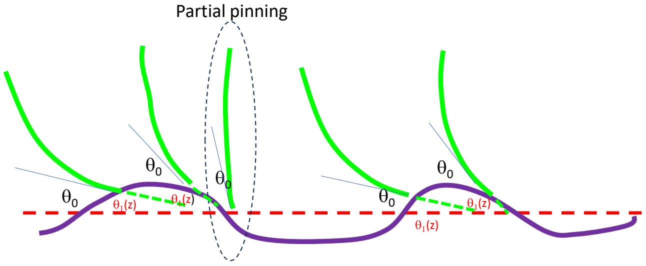

**Figure SI.2.1**. Schematic of the triple line contact on homogeneous rough surface. The capillary pressure is given by the shape of the meniscus (green line), which is approximately linked to the apparent contact angle $\theta_1(z)$ if the roughness relief is small. Each green line represents the fluid front at a given timepoint. The dotted red line represents the position along the axial direction of the channel.

The average apparent contact angle along the length $L$ (in the 1D case) and along the wetted perimeter $p_w$ (in the 3D case) can be defined as

$$\theta_{1,app} = \frac{1}{L}\int_0^L \theta_1(z)\,dz \quad in\ the\ 1D\ case,$$
$$\theta_{1,app} = \frac{1}{L}\frac{1}{p_w}\oint\int_0^L \theta_1(z)\,dzds \quad in\ the\ 3D\ case \qquad \text{(SI2.1)}$$

In the case of a closed channel, it is simply the contact angle for the capillary flow. It is observed that for a homogeneously rough channel, the travel distance vs time approximately follows a square root law of the type Lucas-Washburn as sketched in figure SI.2.2

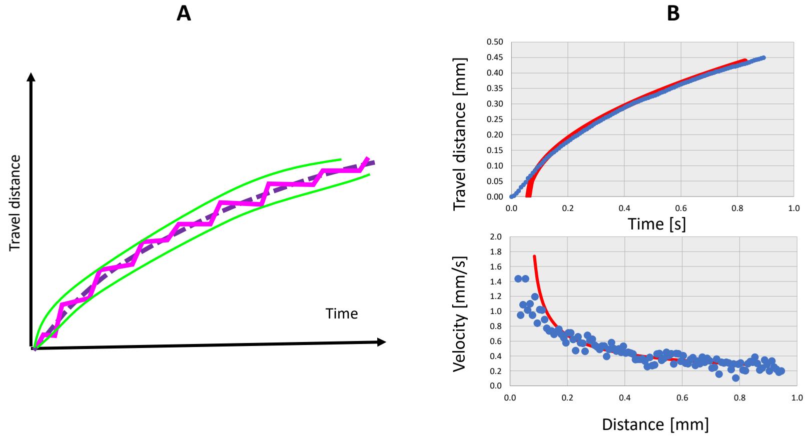

**Figure SI.2.2**. (A) Schematic relating travel distance vs. time for the flow in a rough-walled channel. The purple dotted line shows a general example of observed flow in a 3D printed channel. The alternance of flow pinning and flow restoration occurs on a very short time, so that the "steps" shown by the magenta line are nearly imperceptible. (B) Case of a nonanol flow in the shallow open channel. The red line represents the mLWR model, and the blue dots represent experimental data. Travel distance vs. time is shown in the top graph, and velocity vs. time is shown in the bottom graph.

If at a time $t_1$, the meniscus has reached the length $L_1$, we can write

$$L_1 = \sqrt{\frac{\gamma}{\mu}2\frac{\bar{\lambda}}{\tau}\cos\theta_1^*\ t_1}\ . \qquad \text{(SI2.2)}$$

Then

$$\bar{\lambda}\cos\theta_1^* = \frac{\tau L_1^2}{2t_1}\frac{\mu}{\gamma}, \qquad \text{(SI2.3)}$$

and

$$\cos\theta_1^* = \frac{\tau L_1^2}{2t_1}\frac{\mu}{\gamma}\frac{1}{\bar{\lambda}}\ . \qquad \text{(SI2.4)}$$

Using the definition of the generalized Cassie angle, and for a rectangular open channel

$$cos\,\theta_1^* = \frac{[(w+2h)\;cos\theta_1 - w]}{2\;(w+h)}\,, \qquad \text{(SI2.5)}$$

Then

$$cos\theta_{1,app} = \frac{\left[2\;(w+h)(\frac{\tau L_1^2 \mu 1}{2 t_1 \gamma \bar{\lambda}}) + w\right]}{(w+2h)}\,. \qquad \text{(SI2.6)}$$

And finally

$$\theta_{1,app} = \text{acos}\left\{\frac{\left[2\;(w+h)(\frac{\tau L_1^2 \mu 1}{2 t_1 \gamma \bar{\lambda}}) + w\right]}{(w+2h)}\right\}. \qquad \text{(SI2.7)}$$

**SI.3. Engineering drawings and channel cross section dimension**

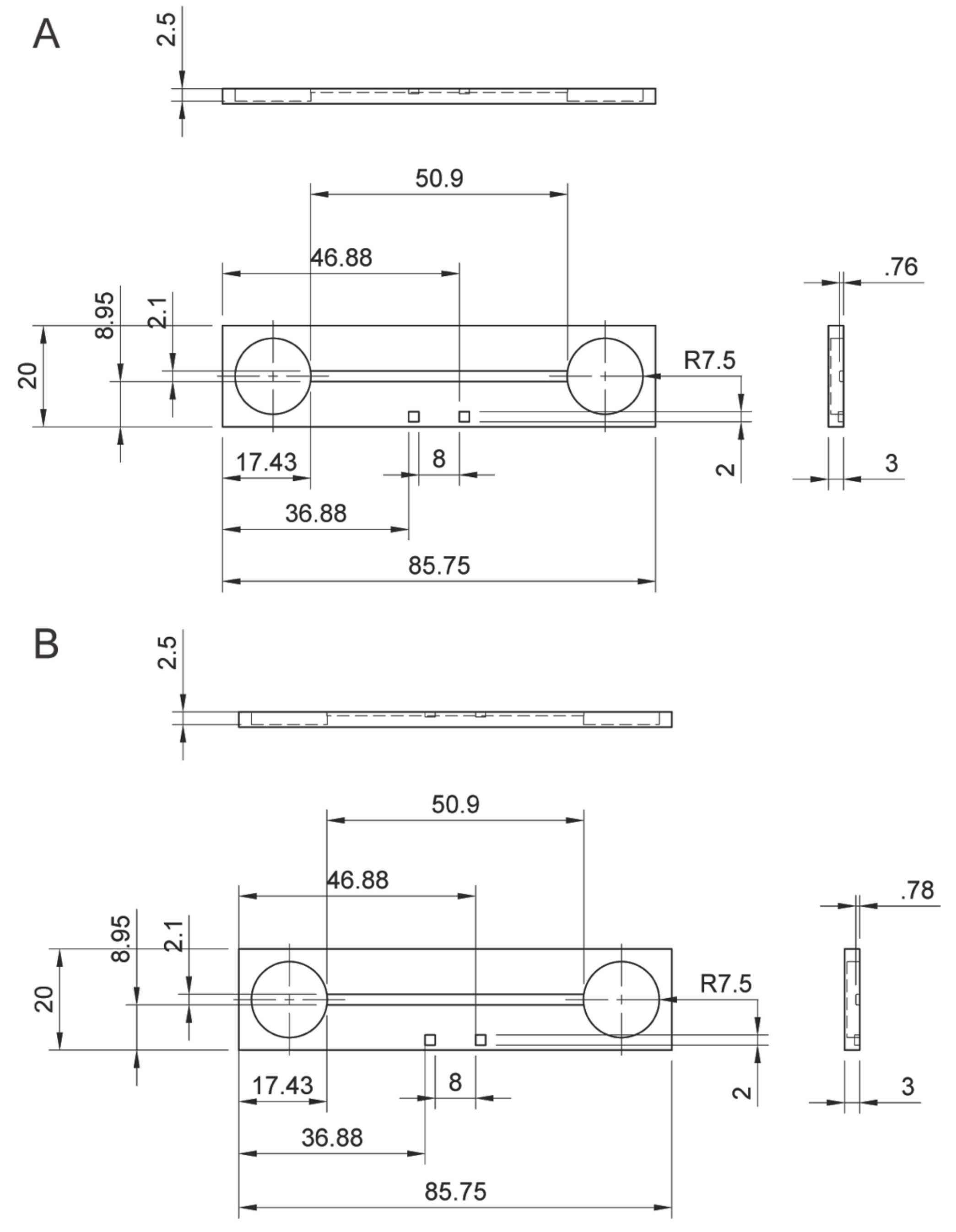

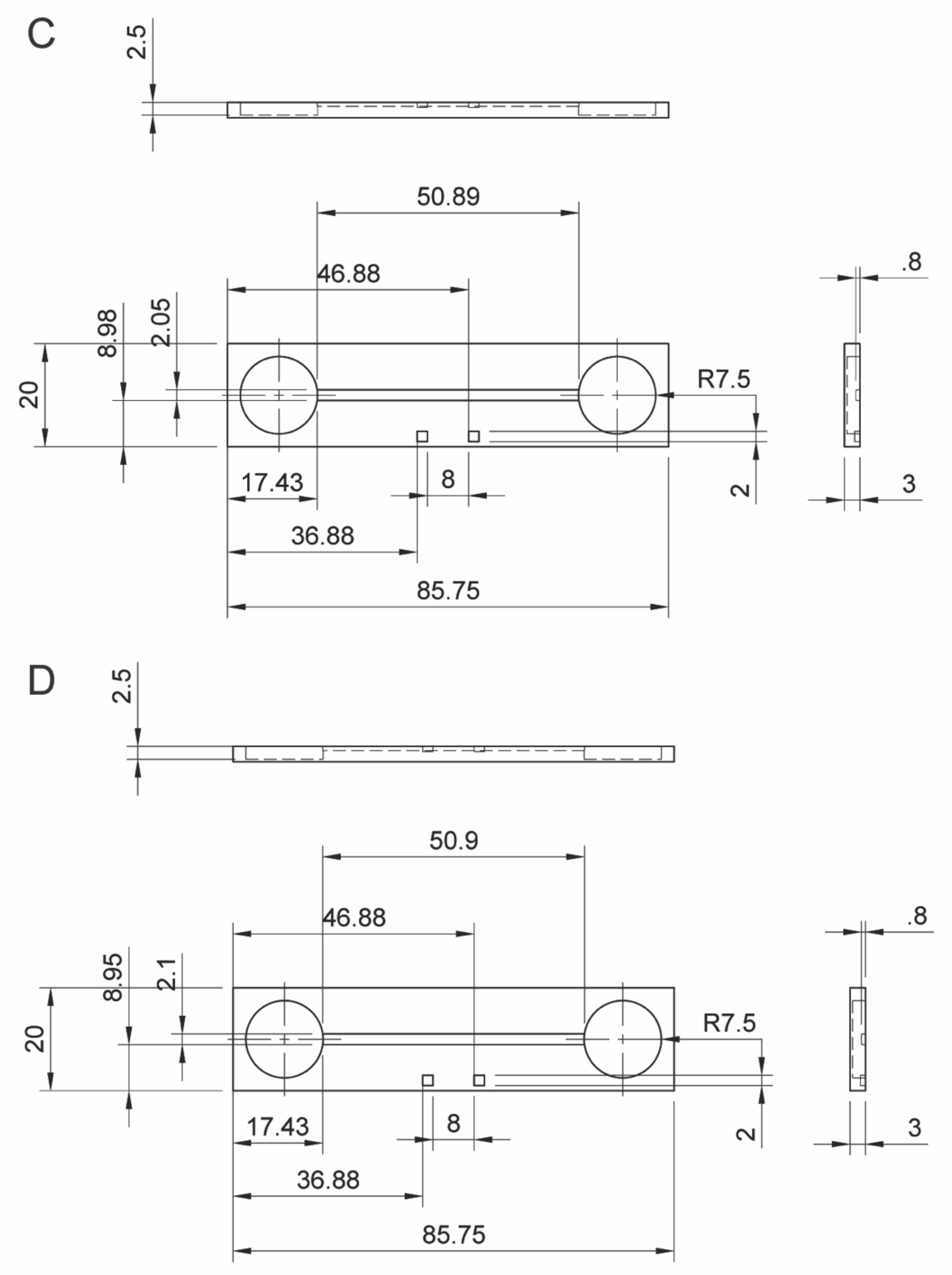
C
2.5
50.89
46.88
.8
8.98
2.05
20
R7.5
17.43
8
2
3
36.88
85.75
D
2.5
50.9
46.88
.8
8.95
2.1
20
R7.5
17.43
8
2
3
36.88
85.75

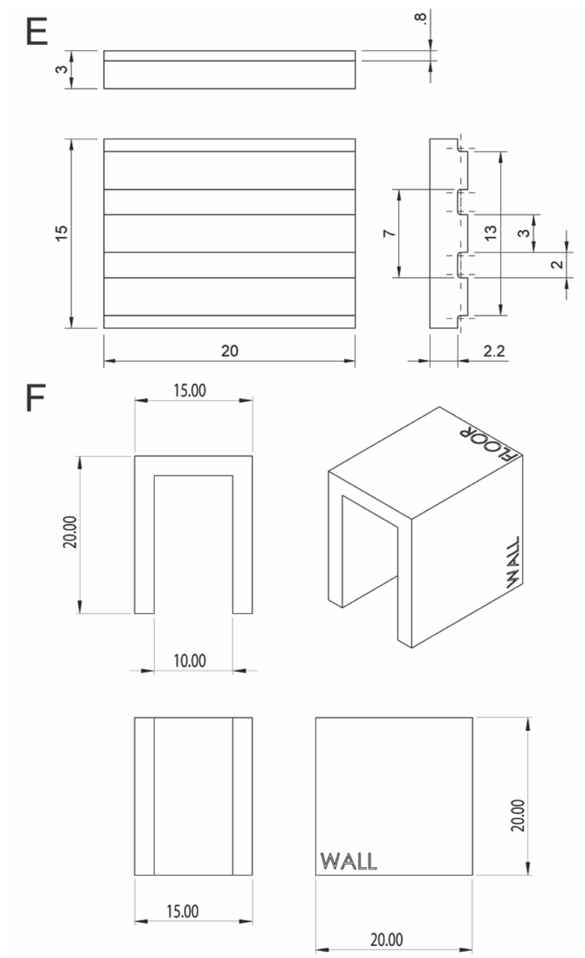

**Figure SI.3.1 Engineering drawings of microfluidic channels.** (A) 0˚ print angle flow device. (B) 15˚ print angle flow device. (C) 45˚ print angle flow device. (D) 90˚ print angle flow device. (E) Representative profilometry block. Channel dimensions are the same across all print angles. (F) Representative contact angle block. Part dimensions are the same across all print angles. All units in mm.

**Table SI.3.1** Designed channel cross section dimensions with respect to print angle to achieve within 5% error of 0.8 mm measured channel height and 2 mm measured channel width.

| | 0˚ print angle | 15˚ print angle | 45˚ print angle | 90˚ print angle |
|---|---|---|---|---|
| Designed channel height | 0.76 mm | 0.78 mm | 0.80 mm | 0.80 mm |
| Designed channel width | 2.10 mm | 2.10 mm | 2.05 mm | 2.10 mm |

### SI.4. Additional experimental data

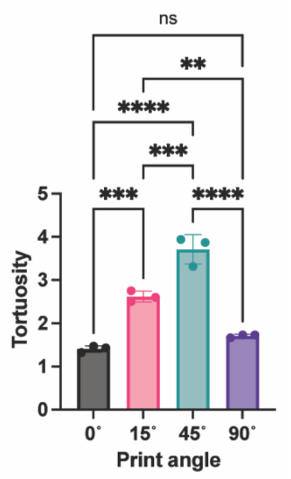

**Figure SI.4.1. Print angle affects surface tortuosity of profilometry block channel floors.** Each point represents the average of 4 measurements on one independently printed part (n=3), and the error bars represent the SD. Statistical analysis was performed using a one-way ANOVA with Tukey's multiple comparisons post hoc test. $^{**}p \leq 0.01$, $^{***}p \leq 0.001$, and $^{****}p \leq 0.0001$.

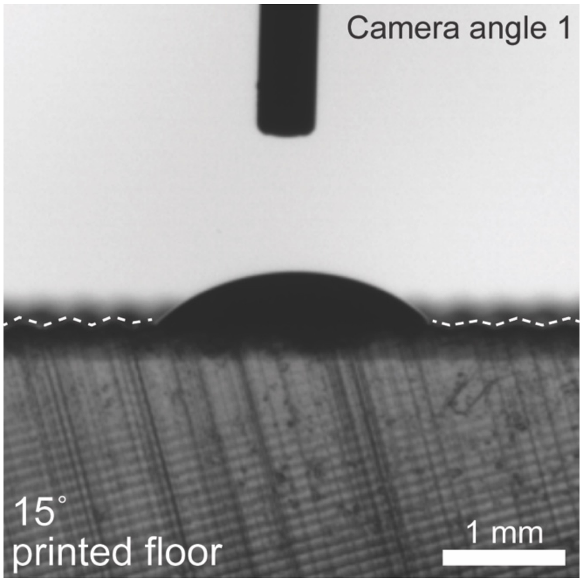

**Figure SI.4.2. Surface texture reliefs created by printing parts at 15˚ are comparable in size to the droplet.** Image of 2 μL droplet of glycerol solution on 15˚ printed contact angle block floor from camera angle 1. The dotted white line follows the outline of the roughness reliefs.

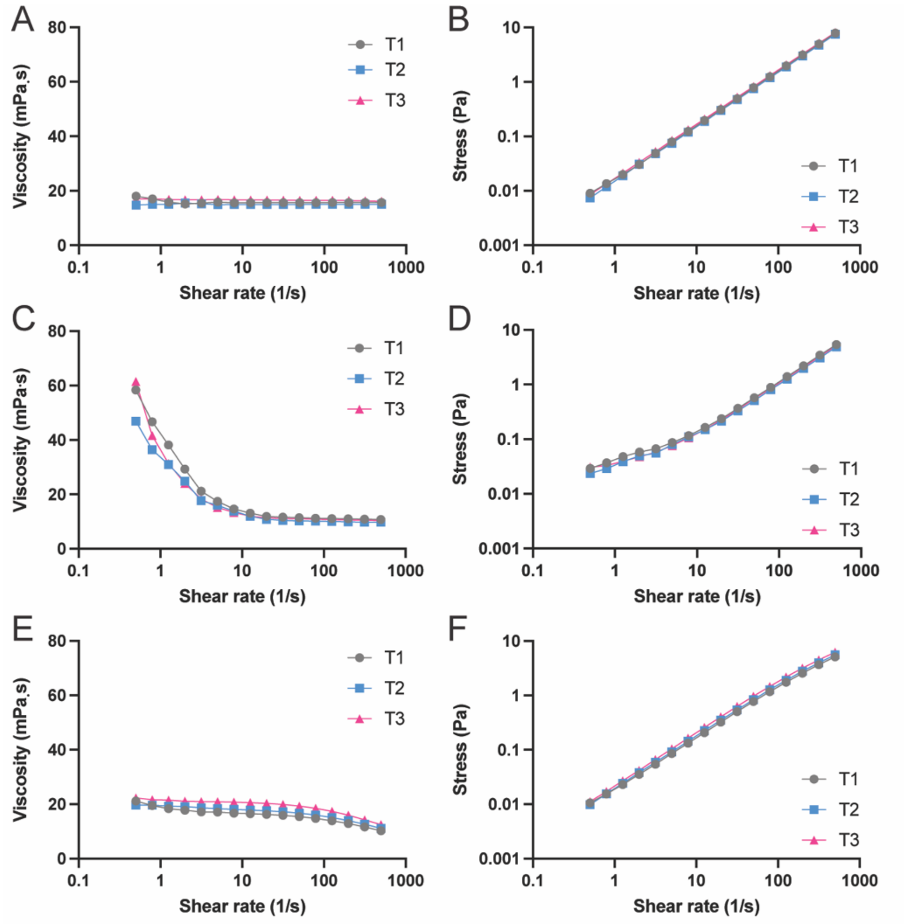

**Figure SI.4.3. Shear rheology of test fluids.** Viscosity and shear stress measurements as a function of shear rate for glycerol solution (A-B), agarose precursor solution (C-D), and collagen precursor solution (E-F). Each line represents an experimental replicate (n=3).

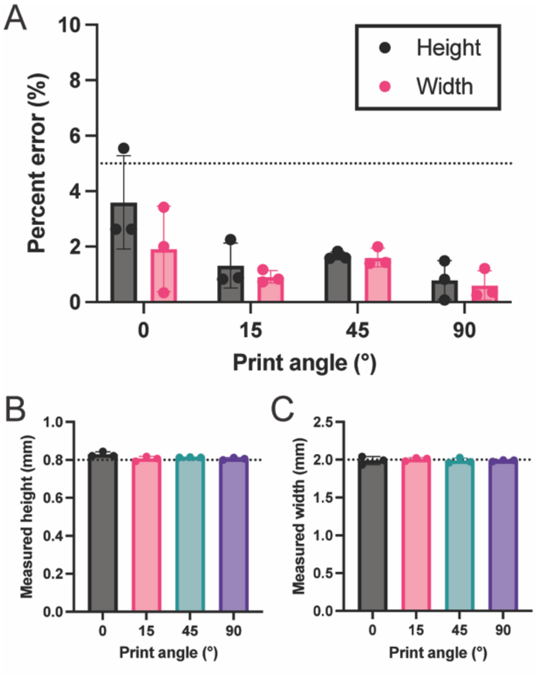

**Figure SI.4.4. Flow device design was reiterated until the measured channel height and width were within 5% error of the target height (0.8 mm) and width (2 mm).** (A) Graph showing percent error of the channel height and width with respect to print angle. (B and C) The measured values of channel height (B) and width (C) are shown for flow devices printed at 0˚, 15˚, 45˚, and 90˚. Each point represents the average of 9 measurements conducted on an independently printed flow device, and error bars represent the SD.

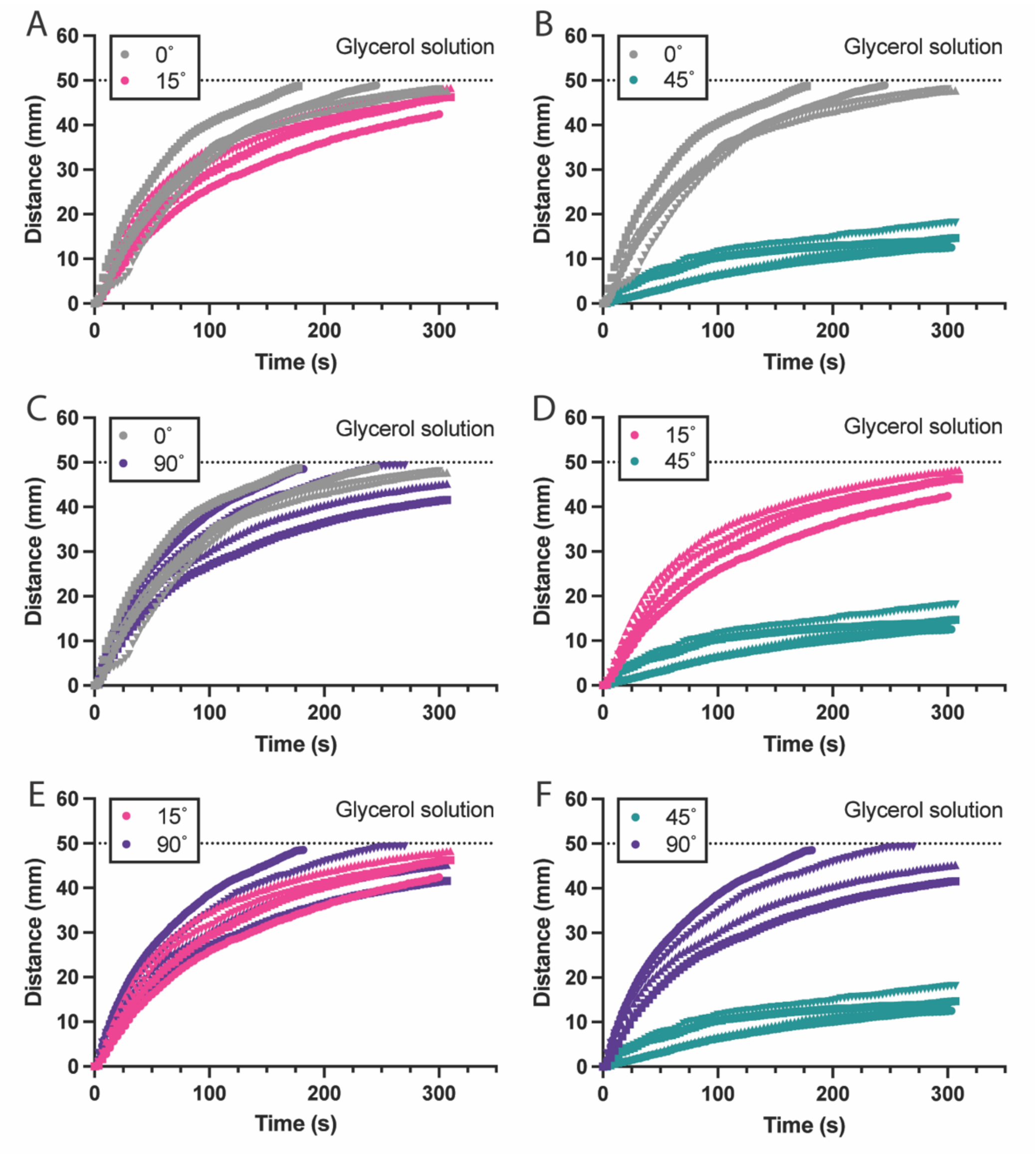

**Figure SI.4.5. Comparison of capillary driven flow with respect to print angle for glycerol solution.** The black dotted lines represent the end of the channel. Each colored line represents flow in an independently printed flow device (n=4).

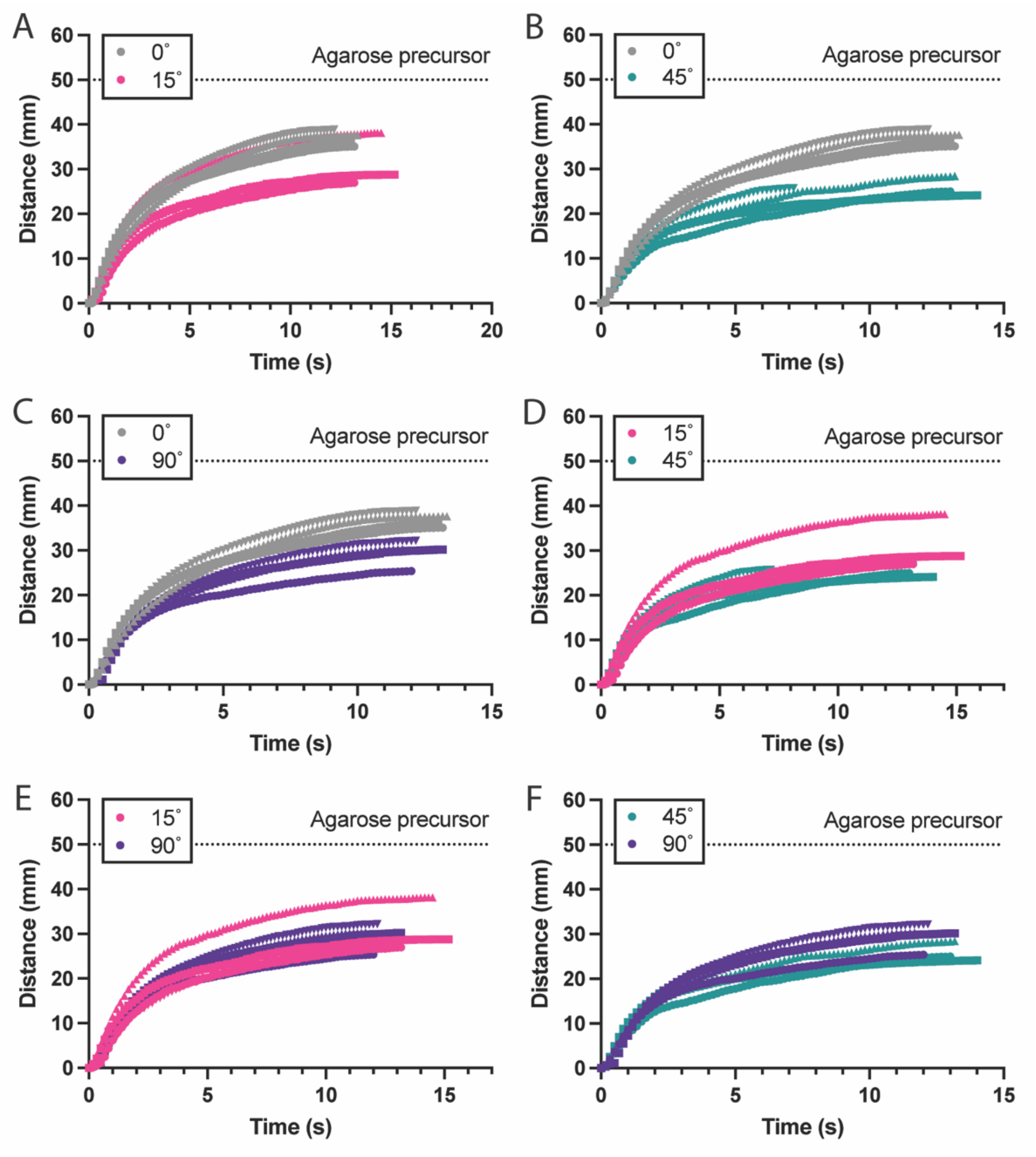

**Figure SI.4.6. Comparison of capillary driven flow with respect to print angle for agarose precursor solution.** The black dotted lines represent the end of the channel. Each colored line represents flow in an independently printed flow device (n=4).

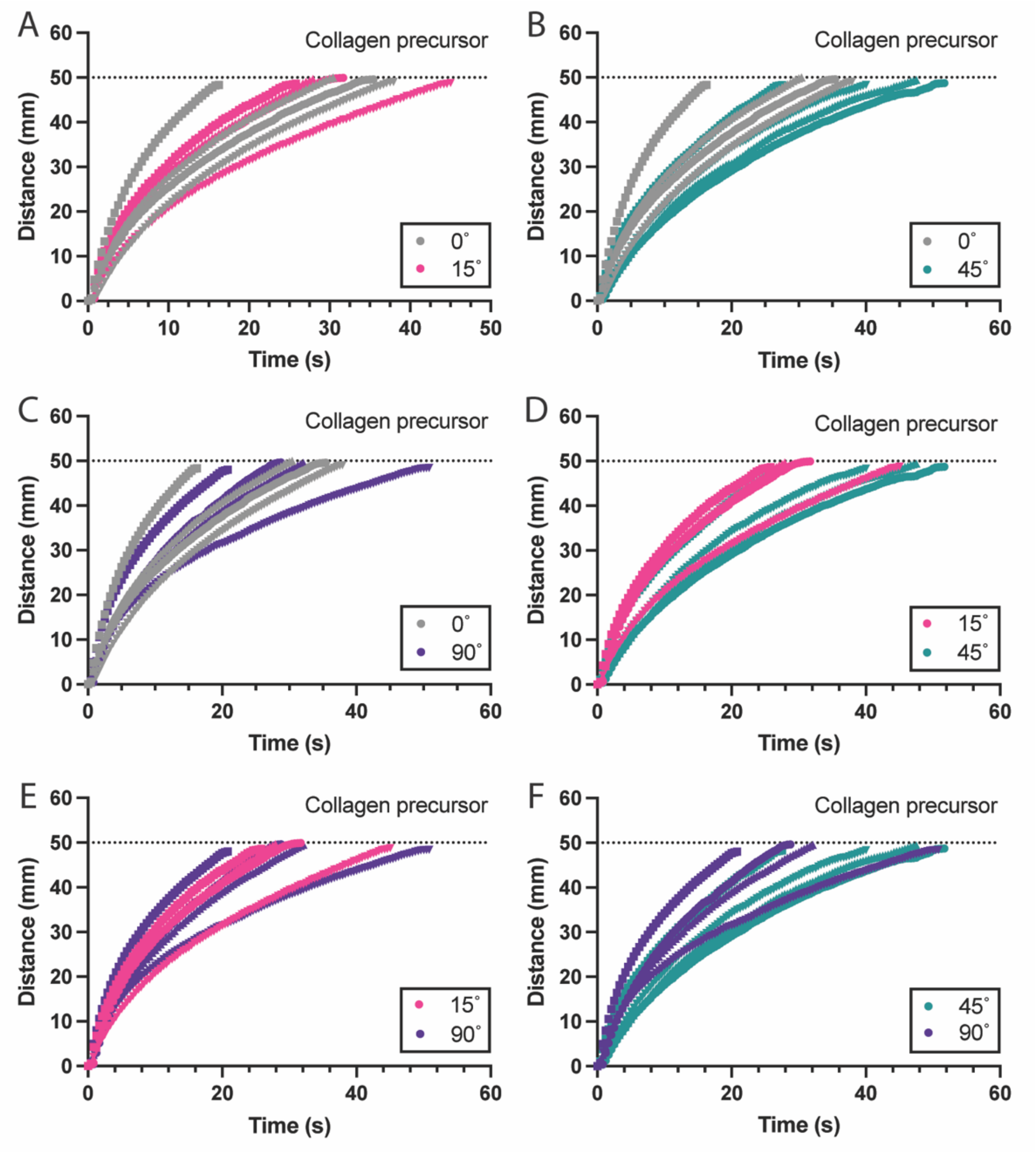

**Figure SI.4.7. Comparison of capillary driven flow with respect to print angle for collagen precursor solution.** The black dotted lines represent the end of the channel. Each colored line represents flow in an independently printed flow device (n=4).

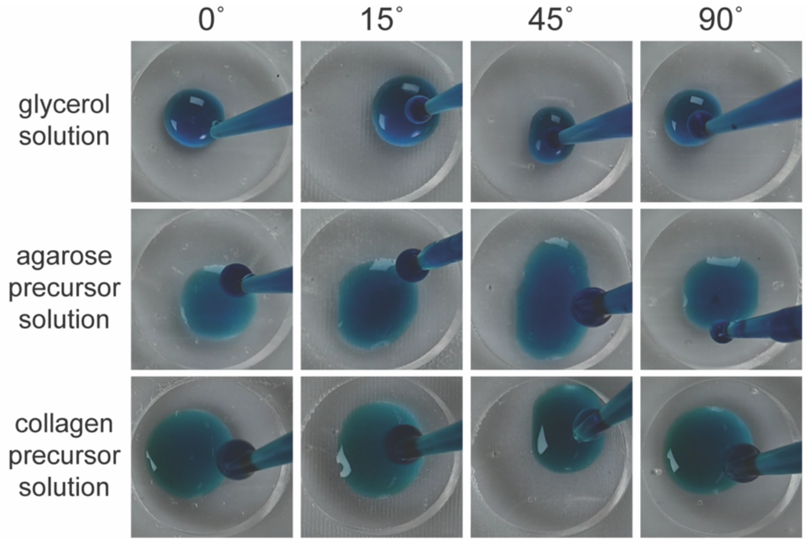

**Figure SI.4.8. Wetting of agarose precursor solution may be greater than indicated by sessile drop measurements.** Images of glycerol solution, agarose precursor solution, and collagen precursor solution when pipetted into flow device reservoirs. Still images were pulled from videos of the first experimental replicate of flow tests. Based on the droplet shape, wetting for agarose precursor solution may be greater than glycerol solution and collagen precursor solution. The drop shape analyzer may not be able to capture this due to the rapid gelling of agarose precursor solution.

### SI.5. Supporting Information References
